# Single-molecule dissection of CFTR folding defects and pharmacological rescue

**DOI:** 10.64898/2026.01.26.701897

**Authors:** Sang Ah Kim, Jesper Levring, Jue Chen, Tae-Young Yoon

**Affiliations:** School of Biological Sciences and Institute for Molecular Biology and Genetics, Seoul National University, Seoul 08826, South Korea; Laboratory of Membrane Biology and Biophysics, The Rockefeller University, New York, NY, USA; Howard Hughes Medical Institute, The Rockefeller University, New York, NY, USA

**Keywords:** ΔF508 CFTR, membrane protein folding, single-molecule magnetic tweezers, pharmacological rescue

## Abstract

Cystic fibrosis is a lethal genetic disorder caused by misfolding of the CFTR protein, most commonly due to the ΔF508 mutation. Despite extensive study, CFTR’s folding process has remained inaccessible to direct observation. Here, we apply single-molecule magnetic tweezers to resolve the complete folding trajectories of wild-type and ΔF508 CFTR with near–amino acid resolution. We find that CFTR follows a hierarchical, template-guided folding pathway in which N-terminal domains scaffold downstream folding. This mechanism tightly couples the free energy states of intermediates, allowing ΔF508-induced instability to propagate across the folding pathway. Pharmacological correctors, in synergy with ATP, reshape the entire folding energy landscape by catalyzing transitions rather than simply stabilizing end states. These long-range, allosteric effects reveal a folding-embedded regulatory network. Our work provides a quantitative framework for mapping multidomain protein folding and therapeutic rescue, offering a broadly applicable strategy for interrogating rare mutations and accelerating structure-based drug discovery.

**Significance Statement:** Misfolding of CFTR underlies cystic fibrosis, and its complex, multidomain architecture makes it an ideal model for understanding how membrane proteins fold and how small molecules can restore native structure. Using single-molecule magnetic tweezers, we reveal how local instabilities propagate through CFTR’s folding pathway and show that pharmacological correctors act by catalyzing specific folding transitions in addition to stabilizing the native fold. These insights establish CFTR as a paradigm for dissecting folding mechanisms in large membrane proteins and for developing general strategies to correct misfolding across diverse human diseases.

## Introduction

The cystic fibrosis transmembrane conductance regulator (CFTR) is a vital member of the ATP-binding cassette (ABC) transporter family uniquely functioning as an anion channel, conducting chloride and bicarbonate across epithelial cell membranes (1, 2). CFTR plays a key role in maintaining the hydration and pH of epithelial secretions, which is critical for physiological function and tissue homeostasis in the lungs, pancreas, intestines, and other organs (3–5).Structurally, CFTR comprises an N-terminal lasso motif, two transmembrane domains (TMDs) that form the anion conduction pore, two cytoplasmic nucleotide-binding domains (NBDs) responsible for ATP binding and hydrolysis (6), and a regulatory (R) domain whose phosphorylation is required for channel activation (7).

Mutations in the *cftr* gene lead to cystic fibrosis, a severe autosomal recessive disorder characterized by salty sweat, chronic respiratory infections, and progressive pulmonary damage (8–10). CFTR mutations are categorized into six classes based on their impact on protein synthesis, folding, gating, and stability (11, 12). The most prevalent disease-causing variant is deletion of phenylalanine at position 508 (ΔF508), which is present in at least one allele in approximately 90% of individuals with cystic fibrosis (13, 14). This mutation disrupts proper protein folding, impairs trafficking to the cell surface, and reduces protein stability, ultimately leading to premature degradation by cellular quality control mechanisms (14–17).

The advent of CFTR modulator therapies has transformed cystic fibrosis treatment by targeting the molecular defects in mutant CFTR proteins. These small-molecule therapies include potentiators, which increase the channel open probability, and correctors, which enhance folding efficiency and promote trafficking of mutant CFTR to the plasma membrane (18–22). Dual-combination regimens of ivacaftor with lumacaftor or tezacaftor, and more recently the triple combination including elexacaftor, have markedly improved clinical outcomes (23). While these therapies now provide substantial clinical benefit to patients with cystic fibrosis, significant challenges remain, particularly for individuals with CFTR mutations that are non-responsive to current modulator therapies, who still lack effective, targeted treatment options (24). Despite extensive efforts to develop treatments for individuals with non-responsive mutations and to address residual dysfunction even in modulator-responsive patients, our understanding of the precise mechanisms by which these modulators act, and the extent to which they effectively rescue CFTR folding and function, remains limited.

CFTR biogenesis has been extensively studied using a variety of methods, including pulse-chase, proteolytic susceptibility assays, antibody mapping, and hydrogen-deuterium exchange mass spectrometry (25–29). These studies have been crucial for tracking the synthesis, maturation and degradation rates of CFTR proteins and collectively supported a two-stage model, consisting of rapid co-translational folding of individual domains and a slower process of post-translational assembly (30, 31). In parallel, structural studies using cryogenic electron microscopy (cryo-EM) have resolved WT CFTR and the ΔF508 variant in distinct conformational states and identified the binding sites of pharmacological molecules (32–38). Despite these progresses, important questions remain regarding the precise kinetics of translation and folding, the nature of intermediate conformational states, the thermodynamic landscape governing folding transitions, and how nascent CFTR is triaged between productive folding and degradation pathways in the context of disease-causing mutations or pharmacological intervention (29, 39, 40). In particular, the dynamic reorganization of interdomain interactions, the mechanisms by which misfolding defects are propagated, and the precise thermodynamic mechanisms by which folding correctors restore native-like conformations remain unresolved.

To bridge this crucial knowledge gap and gain a deeper understanding of CFTR folding dynamics, we employed single-molecule magnetic tweezers (MT). This biophysical technique offers unparalleled resolution in dissecting the folding landscapes of individual protein molecules real-time, an approach previously validated in studies of other complex membrane proteins such as bacterial rhomboid proteases, G-protein-coupled receptors, and glucose transporters (41, 42). Unlike ensemble approaches, single-molecule MT enables direct observation of complete unfolding and refolding trajectories of individual CFTR molecules with near-amino acid resolution, capturing the kinetics and thermodynamics of transient intermediate states that are otherwise inaccessible. By precisely modulating the mechanical force applied to tethered CFTR molecules, we reconstructed the folding and unfolding pathways of both WT and ΔF508 CFTR. This analysis revealed a hierarchical, template-guided folding mechanism in which N-terminal domains fold early and serve as scaffolds to guide the proper folding of downstream regions. Notably, the destabilizing effects of the ΔF508 mutation—and the restorative actions of pharmacological correctors—were found to propagate and amplify along this folding pathway, such that the structural rigidity of early-folding domains directly influenced the folding efficiency of subsequent segments. The quantitative insights gained from this single-molecule approach reveal that distinct combinations of correctors, in conjunction with ATP, actively catalyze folding transitions and function as allosteric modulators that reshape the CFTR folding energy landscape.

## Results

### Reconstruction of full unfolding and refolding processes of CFTR with single-molecule MT

CFTR is a large membrane protein composed of 1,480 residues arranged into five domains with a complex domain-swapping topology (Fig. 1 A and B). We first conducted a series of pilot experiments to test if it is feasible to use single-molecule MT assays to study CFTR unfolding and refolding. To tether CFTR in the magnetic tweezers, we used SpyTag-SpyCatcher system, an engineered peptide-protein pair that forms a spontaneous and irreversible isopeptide bond (43). SpyCatcher was conjugated to the DNA handles and SpyTags were introduced at both termini of the full-length human CFTR (hereafter referred to as WT and ΔF508, denoting the WT and ΔF508 CFTR constructs tagged with SpyTags), enabling stable and site-specific covalent attachment of each handle to the N- and C-termini during force application (SI Appendix, Fig. S1A). Electrophysiological analysis of the dual-SpyTagged CFTR revealed phosphorylation- and ATP-dependent currents typical of CFTR (Fig. 1C) and sensitivity to CFTR modulators, including the potentiator GLPG1837 and the inhibitor CFTR_inh_-172 (SI Appendix, Fig. S1 B and C). Furthermore, the CFTR correctors, lumacaftor and elexacaftor, enhanced the expression of the dual-SpyTagged variant, showing an additive effect similar to that observed with WT CFTR (SI Appendix, Fig. S1D). These findings indicate that the dual-SpyTagged variant preserves the functional and folding properties of WT CFTR and is therefore suitable for MT studies.

**Fig 1.**
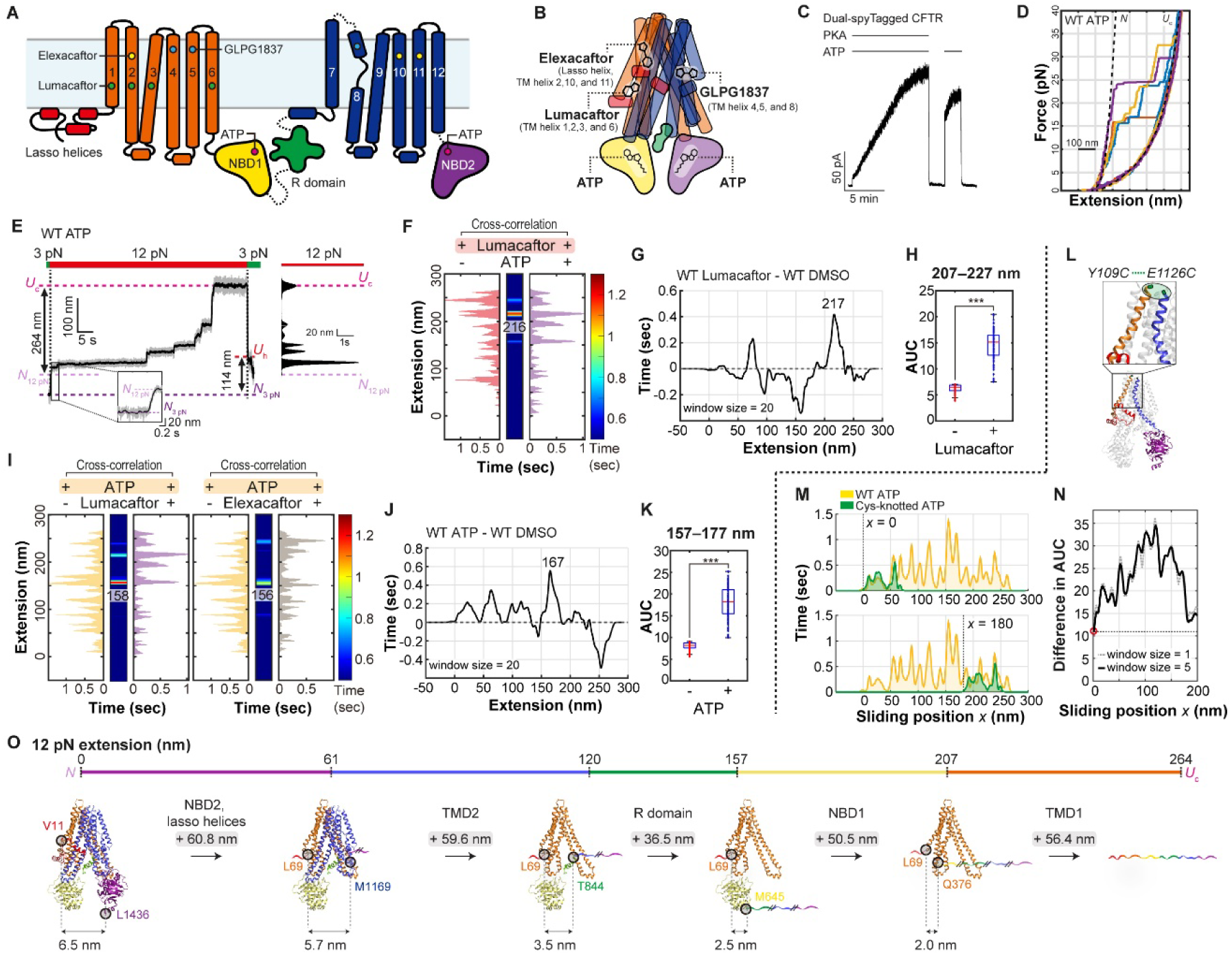
Single-molecule MT assay for CFTR unfolding. (A) Schematic of CFTR domain organization, highlighting the lasso helices (red), TMD1 (orange), NBD1 (yellow), R domain (green), TMD2 (blue), and NBD2 (purple). Individual TM helices are numbered, and the approximate membrane boundaries are indicated. Colored dots indicate the approximate locations of ligand-interacting regions: elexacaftor (yellow), lumacaftor (green), GLPG1837 (blue), and ATP (red) (32). (B) Structural representation of CFTR highlighting the binding sites of small-molecule modulators within the folded tertiary structure of CFTR (32). Lumacaftor binds to a pocket within TMD1, formed primarily by TM helix 1,2,3, and 6. Elexacaftor binds to a distinct composite site involving the lasso helix and elements of TM helix 2, 10, and 11, positioned at the TMD1–TMD2 interface. The potentiator GLPG1837 occupies a hydrophobic cavity formed by TMH4, TMH5, and TMH8, enhancing gating. ATP molecules bind at each NBD1 and NBD2, completing the canonical asymmetric ATP-binding sites. (C) Representative recordings of protein kinase A (PKA)- and ATP-dependent currents from dual-SpyTagged CFTR in inside-out excised patches. 3 mM ATP and 300 nM PKA were used for the recordings. (D) FECs showing unfolding of single CFTR proteins under mechanical tension. Colored FECs represent example traces of individual force-ramp cycles, whereas black dotted lines indicate theoretical FECs for *N* and *U*_c_ states. The experiment was performed using 1 mM ATP. (E) Example force-jump cycle (left) and corresponding histogram at 12 pN (right). Raw data of the exemplary trace at 1.2 kHz are shown in gray and 10-Hz median filtered data in black. (F) Cross-correlation analysis of unfolding histograms with lumacaftor and with or without ATP. The unfolding intermediate at 216 nm extension stabilized by lumacaftor is annotated. The experiment was performed using 1 µM lumacaftor. *n* = 64 (WT Lumacaftor) and *n* = 47 (WT ATP + Lumacaftor) force cycles were used to generate each histogram. (G) Sliding window difference analysis comparing unfolding histograms of WT CFTR in the presence of lumacaftor versus DMSO. A prominent increase in dwell time was observed between 207 and 227□nm, with the peak difference centered at 217□nm. *n* = 64 (WT Lumacaftor) and *n* = 45 (WT DMSO) force cycles were used to generate each histogram prior to subtracting the two histograms with the sliding window. (H) Comparison of AUC in the 207–225 nm extension range with versus without lumacaftor. The boxplot elements represent the following: median (red vertical line); box limits (25^th^ and 75^th^ percentiles); whiskers (extending to 1.5×IQR). Statistical significance was tested using a two-sample t-test (***p ≤ 1 × 10^−3^). (I) Cross-correlation analyses of unfolding histograms with ATP and with or without folding corrector. The unfolding intermediate at approximately 157 nm extension stabilized by ATP is annotated. *n* = 64 (WT ATP), *n* = 47 (WT ATP + Lumacaftor), and *n* = 54 (WT ATP + Elexacaftor) force cycles were used to generate each histogram. (J) Sliding window difference analysis comparing unfolding histograms of WT CFTR in the presence of ATP versus DMSO. A prominent increase in dwell time was observed between 157 and 177□nm, with the peak difference centered at 167□nm. *n* = 54 (WT ATP) and *n* = 45 (WT DMSO) force cycles were used to generate each histogram prior to subtracting the two histograms with the sliding window. (K) Comparison of AUC in the 157–175 nm extension range with versus without ATP. The boxplot elements represent the following: median (red vertical line); box limits (25^th^ and 75^th^ percentiles); whiskers (extending to 1.5×IQR). Statistical significance was tested using a two-sample t-test (***p ≤ 1 × 10^−3^). (L) Molecular structure of CFTR illustrating the engineered disulfide bond between Y109C and E1126C, which restricts unfolding to NBD2 and the lasso helices. The inset depicts a schematic of the disulfide-linked residues. (M) Comparison of unfolding histograms for Cys-knotted CFTR (*n* = 45) and WT CFTR in the presence of ATP. 5-nm sliding window, one-fourth the width used for full-length WT histograms to accommodate the shorter unfolding range of the Cys-knotted, was applied construct and aligned at two alignment positions: *x* = 0 nm (top) and *x* = 180 nm (bottom). (N) Quantification of AUC differences between Cys-knotted and WT CFTR unfolding histograms. The Cys-knotted histogram, averaged with 5-nm (dashed line) and 1-nm (dotted line) sliding windows, was shifted in 1-nm increments across the WT ATP histogram to calculate AUC differences. The minimum difference occurred at *x* = 0 nm, indicating best alignment in the 0–61 nm region, corresponding to unfolding of NBD2 and the lasso helices. (O) Proposed CFTR unfolding pathway. Numbers above unidirectional arrows indicate the extension changes associated with each domain’s unfolding transition. Numbers between bidirectional arrows represent the end-to-end distances of structured regions, with circles marking the N- and C-terminal residues involved in tertiary structure formation.

To prepare CFTR for single-molecule studies, we purified the dual-SpyTagged protein and attached two 512-base-pair DNA handles via SpyTag-SpyCatcher linkages (SI Appendix, Fig. S1 E–H). One DNA handle was anchored to the surface of a polyethylene glycol (PEG)-passivated flow cell, whereas the other was linked to a 3 μm-diameter magnetic bead (SI Appendix, Fig. S2A) (44). Magnetic force was applied to individual CFTR molecules reconstituted in a lipid bicelle environment, cycling gradually between 1 pN and 40 pN (SI Appendix, Fig. S2 A and B). The distance between the two SpyTags, representing the N- and C-termini (hereafter referred to as the extension), was monitored as a function of force (SI Appendix, Fig. S2B). The experimentally obtained force-extension curve (FEC) was compared to theoretical FECs for the native (*N*) and fully unfolded polypeptide (*U*_c_) states of CFTR (Fig. 1D and SI Appendix, Fig. S2 C and D). After repetitive force apply, CFTR proteins extended to yield a FEC matching the predicted trajectory for the *U*_c_ state, lacking any secondary structure at forces exceeding 15 pN (SI Appendix, Fig. S2 C and D; see Methods for extension value calculations). When the force was subsequently reduced to 1 pN and held for a 6-minute incubation, CFTR refolded to the *N* state, evident by the FEC of the subsequent cycle aligning with the theoretical curve of a completely folded CFTR (Fig. 1D).

Encouraged by these observations, we proceeded to establish complete unfolding and refolding pathways under conditions of constant force. In these experiments, we started with CFTR in the *N* state with an initial force of 3 pN, then rapidly increased the force to 12 pN to initiate unfolding (Fig. 1E and SI Appendix, Fig. S2G). With a constant force of 12 pN, the extension of CFTR increased in a stepwise manner, reaching a maximum extension increase of 264 nm, which closely matched the theoretical value for complete unfolding from the *N* to the *U*_c_ state (Fig. 1E and SI Appendix, Fig. S2 E and G). The intermediate extension values observed during this process likely represent progressive unfolding of distinct structural elements. After CFTR reached the *U*_c_ state, we reduced the force to 3 pN to allow refolding while preserving the high-resolution capabilities of single-molecule MT for tracking this process (SI Appendix, Fig. S2 H and I) (41, 42, 45). At 3 pN, the extension of CFTR gradually decreased, with a maximum contraction of 114 nm, precisely matching the expected value for the *N* state (SI Appendix, Fig. S2 F, H and J).

These data demonstrated that the complete unfolding and refolding processes of CFTR could be faithfully reconstituted using our single-molecule MT assay. It was however noteworthy that not all trials resulted in successful refolding. As we will discuss later, the probability of refolding varied between WT and mutant CFTR and was influenced by ligands such as ATP and folding correctors. To gauge the robustness of the assay, we performed blinded experiments in which each trace was acquired in the presence of lumacaftor, elexacaftor, the potentiator GLPG1837, or a DMSO control. DMSO and GLPG1837 produced virtually identical refolding probabilities (∼10 %), whereas lumacaftor and elexacaftor more than doubled this value (>20 %) (SI Appendix, Fig. S3A). The clear separation between correctors and non-correctors highlights the sensitivity of the assay.

### The CFTR unfolding pathway

Unfolding intermediates of WT CFTR were identified by comparing 12-pN unfolding histograms recorded with or without various ligands, with the assumption that each ligand primarily stabilizes the structural region to which it binds. Although each high-force pulling stochastically samples a different set of intermediates at 12 pN, the cumulative histogram converged after inclusion of more than 30 traces (SI Appendix, Fig. S3B). Indeed, 100 independent resampling of 35 traces drawn from the full data set (∼50 traces) produced virtually identical histograms with negligible variance (SI Appendix, Fig. S3 B–E). Through pairwise cross-correlation analyses, we identified intermediates that were elevated in both histograms due to a common ligand (Fig. 1 F, I and SI Appendix, Fig. S3 F–H; see Methods for the cross-correlation analysis). For example, the highest cross-correlation peak between two histograms obtained in the presence of lumacaftor occurred at 216 nm (Fig. 1F and SI Appendix, Fig. S4A), suggesting that lumacaftor stabilizes an intermediate with an extension of approximately 216 nm. Theoretical calculations indicated that the fully extended CFTR at 12 pN reaches 264 nm (SI Appendix, Fig. S2E), with the unfolding of the N-terminal transmembrane domain (TMD1), encompassing transmembrane (TM) helices 1 to 6, alone contributing an extension increase of 57 nm (SI Appendix, Fig. S4B). Thus the 216 nm peak stabilized by lumacaftor aligned closely with the predicted extension of an intermediate where only TMD1 remains folded (SI Appendix, Fig. S4B). In addition, to pinpoint lumacaftor-induced changes in the unfolding profile, we computed sliding-window differences between histograms collected with and without the corrector (Fig. 1G and SI Appendix, Fig. S4 C and D; see Methods for the sliding window difference analysis). We used a 20-nm window—corresponding both to the typical spacing between unfolding histogram peaks and to the expected extension gained upon unzipping a TM-helix hairpin at 12 pN (SI Appendix, Fig. S4D). This analysis revealed a pronounced dwell-time increase centered at 217 nm (Fig. 1G), with significant positive deviations spanning 207–227 nm (Fig. 1H and SI Appendix, Fig. S4E; see Methods for the area under the curve (AUC) analysis). Taken together, these data indicate that TMD1 unfolds last in our MT experiments.

Similarly, cross-correlation analysis indicated that the most prominent peak stabilized by ATP appeared around 157 nm in extension (precisely 158 nm or 156 nm depending on the second ligand) (Fig. 1I and SI Appendix, Fig. S4F). A comparative analysis of the unfolding profiles with and without ATP revealed a marked increase in dwell time (Fig. 1J and SI Appendix, Fig. S4G), with significant positive differences spanning the 157–177 nm region (Fig. 1K). Furthermore, the presence of ATP notably enhanced sampling in earlier extension regions: in particular, the extension region from 0 to 61□nm exhibited virtually no sampling in the absence of ATP (SI Appendix, Fig. S4G). Given that ATP binds to both NBD1 and NBD2 (Fig. 1B and SI Appendix, Fig. S5A), our observations suggest that the 157–177 nm region most likely corresponds to NBD1, which harbors the degenerate ATP-binding site defective of hydrolyzing ATP (46). We further hypothesize that the extension increase from 0 to 61 nm reflects unfolding of NBD2 and the lasso helices—an assignment that aligns with the pulling geometry of our MT assay, where unfolding is likely to initiate from either the N- or C-terminus.

To test this hypothesis, we first engineered a CFTR variant containing cysteine substitutions at Y109 and E1126 (Fig. 1L and SI Appendix, Fig. S1 I–M). This modification introduced a ‘knotted’ topology that restricted unfolding to NBD2 and the lasso helices in the absence of the reducing agents (Fig. 1L and SI Appendix, Fig. S5B). The unfolding histogram of the ‘knotted’ CFTR was compared to sliding windows of the WT CFTR unfolding histogram (in the presence of ATP) along the extension axis (Fig. 1M). This comparison revealed that the smallest difference between the histograms occurred when the WT CFTR signal was aligned to the 0–61 nm extension range (Fig. 1 M, N and SI Appendix, Fig. S5C). In a second test, we disrupted ATP binding to NBD2 by introducing the Y1219A mutation, which eliminates stacking interactions at the ATP site (SI Appendix, Fig. S5A). While ATP still enhanced the 156 nm peak (SI Appendix, Fig. S5 D and E), the Y1219A mutation significantly reduced dwell times in the 0–61 nm region (SI Appendix, Fig. S5 F–H). Together, these results support our hypothesis that the initial 61 nm extension arises from unfolding of NBD2 and the lasso helices (SI Appendix, Fig. S5I), while the extension increase from 157 to 206 nm predominantly reflects NBD1 unfolding (SI Appendix, Fig. S5J).

Finally, we identified a consistent valley in the unfolding histograms, spanning from 120 to 140 nm across all reaction conditions tested (SI Appendix, Fig. S5K). The absence of significant peaks in this region suggests the unfolding of a disordered segment, permitting us to infer that this region corresponds to the extension of CFTR’s R domain.

Together, these data allowed us to construct a detailed unfolding pathway for CFTR (Fig. 1O). Under a mechanical force of 12 pN, CFTR unfolds in a domain-specific sequence. The process begins with the unfolding of the N-terminal lasso helices and the C-terminal NBD2, followed by the six TM helices in TMD2. This is then succeeded by the unfolding of the R domain, NBD1, and finally TMD1. Remarkably, ligands can bind to CFTR even when suspended between magnetic tweezers, stabilizing specific domains from unfolding. These findings demonstrate that ligands such as ATP and lumacaftor can directly enhance the structural stability of their respective binding domains.

### The CFTR refolding pathway

We next studied the *in vitro* CFTR refolding pathway at a single-molecule level. As described earlier, following the unraveling of each CFTR molecule to the fully extended *U*_c_ state, the magnetic force was reduced and held at 3 pN to allow CFTR refolding (Fig. 2A and SI Appendix, Fig. S6A). Based on the measured 114 nm extension during refolding (Fig. 2A) and our theoretical calculation (SI Appendix, Fig. S2F), we propose that upon force relaxation to 3 pN, CFTR rapidly transitions into an unfolded helical (*U*_h_) state, where only the helical secondary structures are restored (41, 42). The formation of tertiary structures follows over the course of minutes, with only a 10% probability of achieving the fully folded native (*N*_3pN_) state for the WT CFTR without ligands (Fig. 2A, inset). The addition of lumacaftor or ATP substantially increased the refolding probability to 26% and 39%, respectively (Fig. 2A, inset).

**Fig 2.**
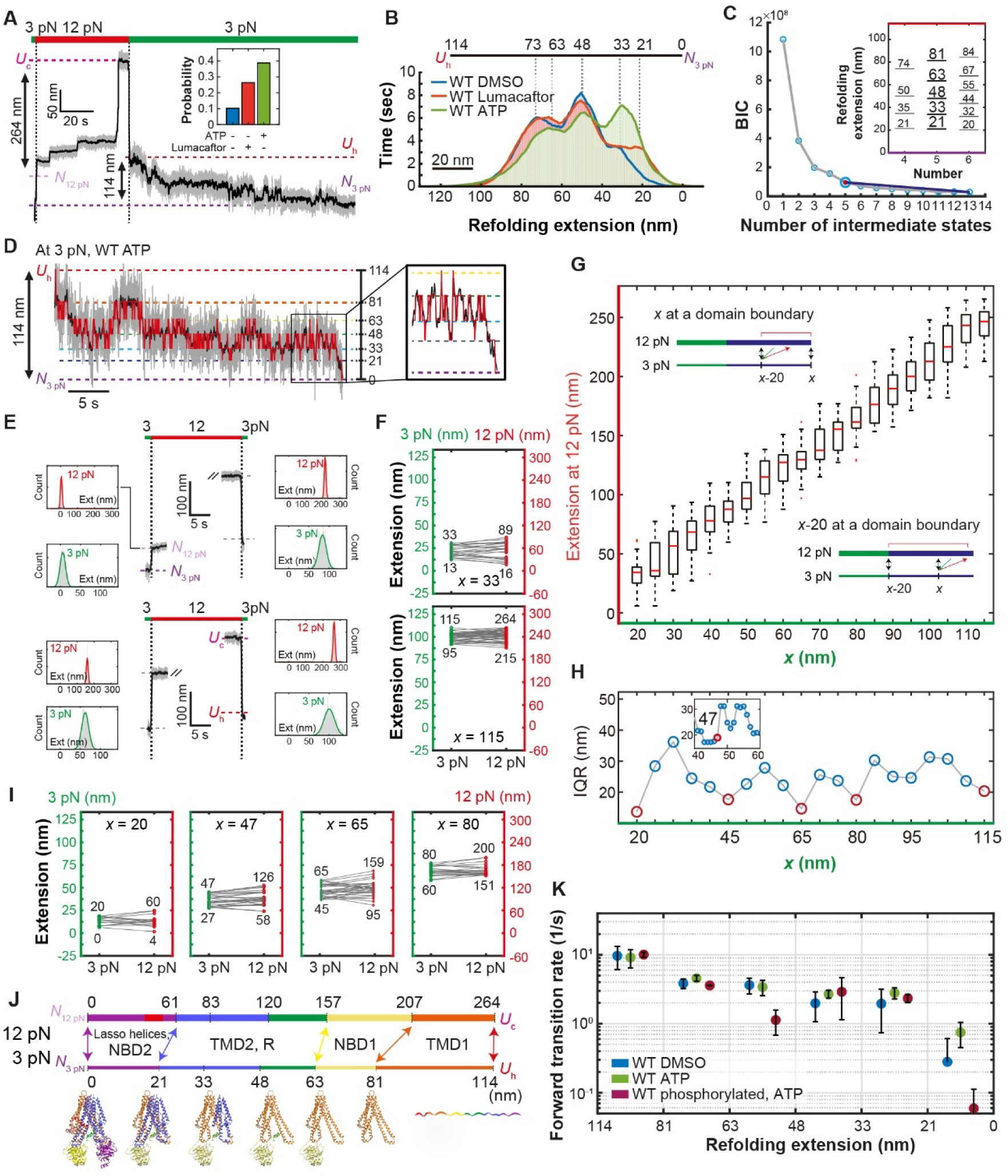
Single-molecule MT assay for CFTR refolding. (A) Experimental protocol for studying CFTR refolding dynamics. Single CFTR proteins were held at 3 pN after relaxation from 12 pN to monitor refolding. Inset shows the probability of complete refolding; the first two data points are replotted from SI Appendix, Fig. S3A. (B) Refolding histograms generated with a 10-Hz median filter reveal multiple distinct peaks, corresponding to refolding intermediates. *n* = 27 (WT DMSO), *n* = 34 (WT ATP), and *n* = 32 (WT Lumacaftor) refolding traces were used to generate each histogram. (C) BIC analysis identifies five as the optimal number of intermediate states in WT CFTR refolding traces. The inset shows the positions of intermediate states for an assumed number of states. The analysis was performed using WT ATP. (D) Representative refolding trace with 1.2 kHz raw data (gray) and 10-Hz median filtered data (black). HMM analysis was applied to refolding traces and the Viterbi path was overlaid on the data (red). Horizontal lines indicate extension values shown on the right. (E) Force-jump experiments to record extension states at 3 pN before and after a jump to 12 pN. Gaussian fitting was applied to measure precise mean extension values before and after the force change. (F) Examples of mapping corresponding extension values between 3 pN and 12 pN from force-jump experiments. Numbers below and above the green and red circles indicate the minimum and maximum extensions observed at 3 pN and 12 pN, respectively, for a given *x*. (G) Correlative mapping between 3-pN and 12-pN extension spaces based on force-jump experiment data. Extension data were grouped into 20-nm windows, with *x* denoting the upper boundary of the window in the 3-pN extension space. Box plots depict the median (red line), IQR (box limits), and 1.5×IQR (whiskers). Data reflect *n* = 164 force-jump traces. The inset shows reduced variance occurs when *x* corresponds to a domain boundary (see SI Appendix, Fig. S6 G and H for detailed explanation). (H) IQR for different values of *x*. Local minima correspond to distinct structural regions at 20, 65, 80 and 115 nm reflecting domain boundaries. Around *x* = 45 nm, detailed IQR analysis at 1-nm resolution uncovers a sharp increase in IQR above 47 nm (inset), indicating a structural transition point. (I) Mapping the exact extension values at 12 pN for the local minima identified in Fig. 2H. (J) Correlative comparison of unfolding intermediates at 12 pN and refolding intermediates at 3 pN demonstrates that refolding proceeds in the exact reverse sequence of unfolding, consistent with a templated folding pathway. (K) Forward transition rates among refolding intermediates for non-phosphorylated CFTR in the presence of DMSO or ATP and for phosphorylated CFTR in the presence of ATP. 300 nM PKA was used to phosphorylated CFTR. Data represent means and standard deviations for *n* individual traces. *n* = 27 for WT DMSO, *n* = 34 for WT ATP and *n* = 24 for WT phosphorylated, ATP.

Refolding histograms, generated by compiling all the extension values sampled during these 3 pN refolding trials (with 10-Hz median filtering applied) displayed several peaks, indicative of distinct refolding intermediates (Fig. 2B and SI Appendix, Fig. S6B). To independently determine the positions of refolding intermediates, we analyzed individual refolding traces with a hidden Markov model (HMM) (Fig. 2 C, D and SI Appendix, Fig. S6A) (47, 48). Using the Bayesian Information Criterion (BIC), we identified five most likely intermediates, with extension values of 21, 33, 48, 63, and 81 nm, respectively (Fig. 2C, inset). These values closely corresponded to the peaks observed in the refolding histograms (Fig. 2 B, C and SI Appendix, Fig. S6B).

To explore the relationship between the 3 pN refolding intermediate states and the unfolding intermediates observed at 12 pN, we performed a force-jump experiment, rapidly alternating the applied force between 3 and 12 pN. We recorded the extension states immediately before and after each force jump, regardless of the jump direction (Fig. 2E and SI Appendix, Fig. S6 C–E). The extension values at the two force levels showed a linear relationship, validating the force-jump method (SI Appendix, Fig. S6F). We then grouped the 3-pN extension states into 20-nm windows, with the upper limit of each window defined as *x* (e.g., *x* = 33 nm for a window spanning 13–33 nm) (Fig. 2F). Consistent with the linear correlation, the 20-nm windows at 3 pN corresponded closely to distinct clusters at 12 pN (Fig. 2 F and G). An intriguing observation is that the broadness of the 12-pN extension clusters, as measured by the interquartile range (IQR), varied periodically (Fig. 2H). The local minima, at *x* values of 20, 47, 65, and 80 nm, align closely with the refolding intermediates identified at 21, 48, 63, and 81 nm (Fig. 2H versus Fig. 2 B and C). This alignment led us to hypothesize that the minimal spread at 12 pN occurred when *x* corresponds to a domain boundary (Fig. 2G, insets), as fully folded domains are more stable thereby exhibiting minimal structural changes during force jumps (SI Appendix, Fig. S6 G and H).

Furthermore, the mapping between 3 and 12 pN extensions along both directions of force-jump was asymmetric, extending into larger 12 pN values, rather than extending into the smaller 12 pN values (SI Appendix, Fig. S6 I and J). This observation supports the notion that the upper boundary of each 20-nm window at *x*, rather than the lower at *x*-20 nm, should align with domain boundaries, resulting in a tighter IQR (Fig. 2G, insets and SI Appendix, Fig. S6 G and H). Finally, analyzing the upper limits of the 12-pN extension clusters—indicative of domain boundaries—revealed positions at 60, 126, 159, 200, and 264 nm at 12 pN (Fig. 2 F and I, upper value of 12-pN extension). These values closely matched domain boundaries determined in high-force unfolding trials (Fig. 1O and 2J), suggesting that folding occurs in the reverse order of unfolding.

For structural identification of the refolding intermediate at 48 nm, which is a prominent peak in the refolding histogram, we compared the refolding kinetics of dephosphorylated CFTR versus phosphorylated CFTR (Fig. 2K). The transition rates between intermediates, determined from the dwell times at each intermediate, progressively decreases as the protein refolds, with the 21-to-0 nm transition—representing NBD2 and lasso helices folding—being the slowest step (Fig. 2K). The early stages of folding—specifically transition from 114 to 81 nm—occurred rapidly, which accounts for the lack of a discernible 81 nm peak in the refolding histograms (Fig. 2 B, K and SI Appendix, Fig. S6B). Phosphorylation of the R domain selectively altered the later stages of refolding, affecting transitions between 63 and 48 nm, and between 21 and 0 nm (Fig. 2K). Analyzing multiple possible scenarios (SI Appendix, Fig. S7 A and B) indicate that the 48 nm intermediate most likely represents an intermediate with the R domain assembled onto TMD1/NBD1 (SI Appendix, Fig. S7B, scenario 2-2). Supporting this hypothesis, cryo-EM studies indicated that the dephosphorylated R domain, although largely lacking defined secondary structures, packs along the inner surface of NBD1 (32).

Lastly, the observation that TMD2 completed its folded structure at the 21 nm extension led us to characterize the conformational state at 33 nm, where TMD2 appeared to be partially assembled, extending up to half of TM heli× 10 (Fig. 2J and SI Appendix, Fig. S7B, scenario 2-2). It is particularly notable that the transition from 33 to 21 nm encompasses the completion of TMD2, including the formation of the TM helices 11 and 12 and its association with NBD1.

Collectively, these results provide an *in vitro* refolding pathway of CFTR, which closely mirrors that of the unfolding process (Fig. 2J). TMD1 folds first and rapidly, followed sequentially by NBD1, the R domain, TMD2, and finally, NBD2 along with the N-terminal lasso helices. The overall process has a low success rate, which is improved by the folding corrector lumacaftor and ATP, likely through the stabilization of TMD1 and the NBDs, respectively (Fig. 2A, inset).

### Unfolding and refolding pathways of **Δ**F508 CFTR

In the cellular environment, the ΔF508 variant exhibits severe folding and trafficking defects that underlie the pathogenesis of cystic fibrosis (49–51). Cryo-EM structure of ΔF508 CFTR reveals that ΔF508 NBD1 is too flexible to stably associate with the TM helices (36). This defect further disrupts proper positioning of the R domain, as it packs onto the inner surface of NBD1 (36). We next asked whether these structural defects alter the unfolding and refolding pathways, creating alternative routes through a distorted folding energy landscape.

To measure the altered folding properties of the ΔF508 variant, we expressed and purified the dual-SpyTagged ΔF508 CFTR (Fig. 3A and SI Appendix, Fig. S8 A and B). Expression of the ΔF508 variant required inclusion of lumacaftor and elexacaftor in the culture medium. After purification, the folding correctors were removed by dialysis as assessed by high pressure liquid chromatography (HPLC) (SI Appendix, Fig. S8 C and D) and extensive washing steps were conducted to remove residual folding correctors during the single-molecule MT experiment. Using the same force-ramp cycles, we observed that ΔF508 CFTR began unfolding at much lower force levels compared to WT CFTR, indicating reduced stability (compare Fig. 1D with Fig. 3B). Despite this, ΔF508 CFTR displayed distinct unfolding intermediates very similar to those of the WT CFTR (Fig. 3C). Moreover, during unfolding under a constant 12 pN force, the addition of lumacaftor consistently resulted in a prominent cross-correlation peak at 211 nm (Fig. 3D and SI Appendix, Fig. S9A). Comparative analysis of the unfolding histograms with and without lumacaftor revealed that the dwell time in the extension region spanning 207–227 nm was prominently increased by lumacaftor (Fig. 3 E, F and SI Appendix, Fig. S9B). Thus, similar to WT CFTR, the final 57 nm extension observed during the unfolding of ΔF508 CFTR is likely attributable to the unfolding of TMD1 (SI Appendix, Fig. S4B). However, unlike WT CFTR, ATP did not enhance the unfolding peaks corresponding to NBD1 (i.e., between 157 and 206 nm extension), but instead caused a slight decrease (Fig. 3 G–I and SI Appendix, Fig. S9 C and D). These findings, consistent with prior studies (36, 52), indicate that the ΔF508 NBD1 was too flexible to be stabilized by ATP.

**Fig 3.**
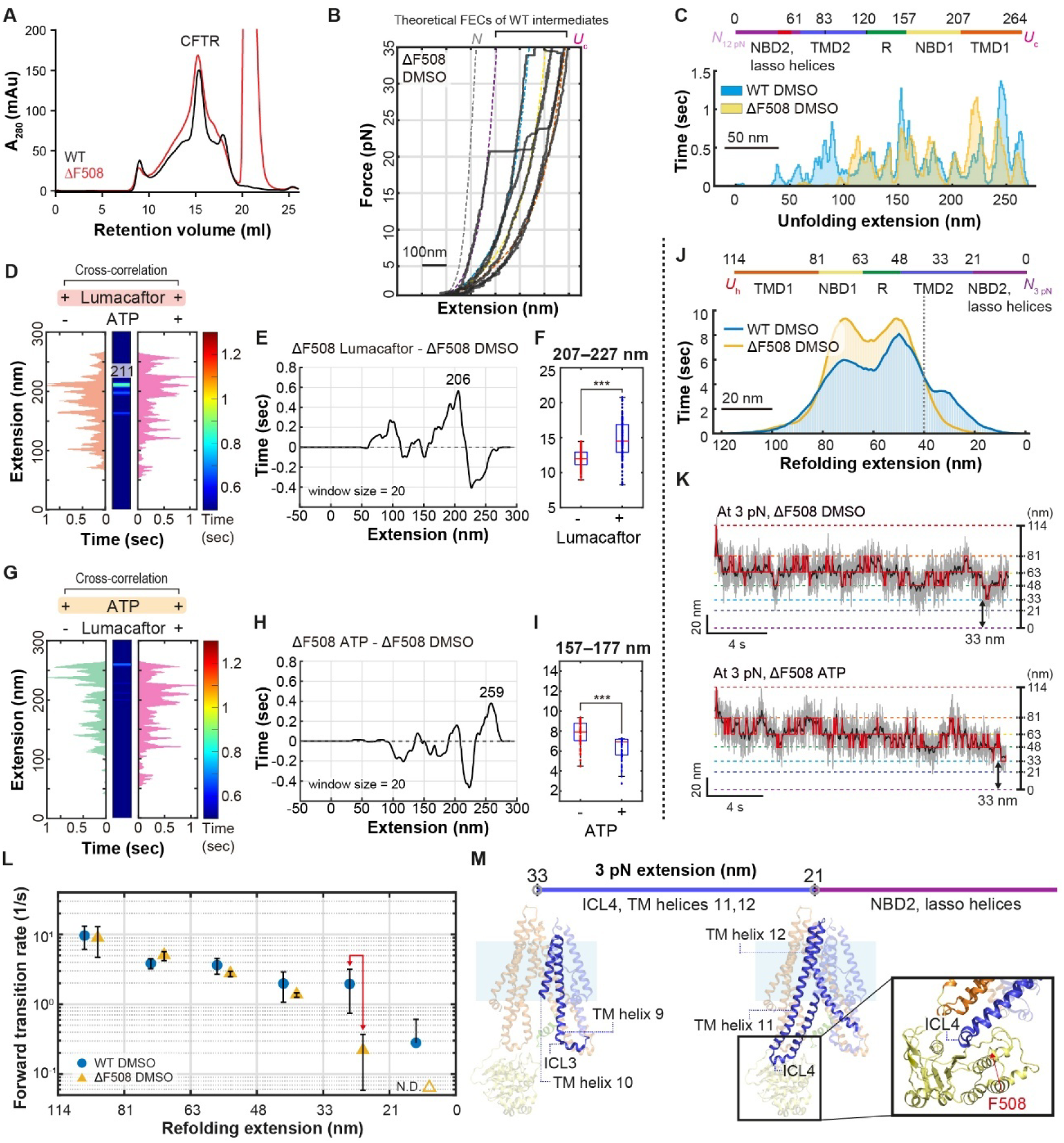
Single-molecule unfolding and refolding of ΔF508 CFTR. (A) Gel filtration profiles of purified digitonin-solubilized WT and ΔF508 dual-SpyTagged CFTR. The WT profile is replotted from SI Appendix, Fig. S1E. (B) FECs from repeated force-ramp cycles of ΔF508 DMSO. Colored dotted lines represent theoretical FECs for individual intermediate states of WT CFTR estimated based on the extensible Worm-Like Chain (eWLC) model. The color code for the theoretical FECs is as follows: gray (native), purple (NBD2 and the lasso helices unfolded), blue (NBD2, lasso helices and TMD2 unfolded), yellow (NBD2, lasso helices, TMD2, R and NBD1 unfolded), and orange (all domains unfolded). (C) Unfolding histograms for WT and ΔF508 CFTR in the absence of ligands but with DMSO. *n* = 48 (ΔF508 DMSO) force cycles were used to generate the histogram. The unfolding histogram for WT DMSO is replotted from SI Appendix, Fig. S4G. (D) Cross-correlation analysis reveals a prominent peak at 211 nm extension in the presence of lumacaftor, with or without ATP. *n* = 70 (ΔF508 Lumacaftor) and *n* = 50 (ΔF508 ATP + lumacaftor) force cycles were used to generate each histogram. (E) Sliding window difference analysis comparing unfolding histograms of ΔF508 CFTR in the presence of lumacaftor versus DMSO. A prominent increase in dwell time was observed between 196 and 216□nm, with the peak difference centered at 206□nm. *n* = 70 (ΔF508 Lumacaftor) and *n* = 48 (ΔF508 DMSO) force cycles were used to generate each histogram prior to subtracting the two histograms with the sliding window. (F) Comparison of AUC in the 207–227 nm extension range (corresponding to the same range used for WT CFTR), with versus without lumacaftor. The boxplot elements represent the following: median (red vertical line); box limits (25^th^ and 75^th^ percentiles); whiskers (extending to 1.5×IQR). Statistical significance was tested using a two-sample t-test (***p ≤ 1 × 10^−3^). (G) Cross-correlation analysis reveals no prominent peak at the extension corresponding to NBD1 in the presence of ATP, with or without lumacaftor. *n* = 45 (ΔF508 ATP) and *n* = 50 (ΔF508 ATP + Lumacaftor) force cycles were used to generate the histogram. (H) Sliding window difference analysis comparing unfolding histograms of ΔF508 CFTR in the presence of ATP versus DMSO. No prominent increase in dwell time was observed between 157 and 177□nm. *n* = 45 (ΔF508 ATP) and *n* = 48 (ΔF508 DMSO) force cycles were used to generate each histogram prior to subtracting the two histograms with the sliding window. (I) Comparison of AUC in the 157–177 nm extension range with versus without ATP. The boxplot elements represent the following: median (red vertical line); box limits (25^th^ and 75^th^ percentiles); whiskers (extending to 1.5×IQR). Statistical significance was tested using a two-sample t-test (***p ≤ 1 × 10^−3^). (J) Refolding histograms at 3 pN for WT and ΔF508 CFTR in the absence of ligands but with DMSO. *n* = 27 (WT DMSO) and *n* = 27 (ΔF508 DMSO) refolding traces were used to generate each histogram. The refolding histogram for WT DMSO is replotted from Fig. 2B. (K) Representative refolding trace with 1.2 kHz raw data (gray) and 10-Hz median filtered data (black). HMM analyses were applied to refolding traces and the Viterbi path was overlaid on the data (red). Horizontal lines indicate extension values shown on the right. (L) Forward rate constants for folding transitions of WT and ΔF508 CFTR with DMSO. The red arrow highlights the effect of the ΔF508 mutation on the transition from 33 to 21 nm. Data represent means and standard deviations for *n* individual traces. *n* = 27 for ΔF508 DMSO. The transition rates for WT DMSO are replotted from Fig. 2K. N.D., not determined. (M) Structural model of CFTR folding intermediates corresponding to extension values of 33 nm and 21 nm at 3 pN.

When the force was reduced to 3 pN, ΔF508 CFTR, like WT CFTR, began to refold in a stepwise manner (Fig. 3 J and K). While compiled refolding histograms and HMM analyses of individual refolding traces identified intermediates closely resembling those of WT CFTR, refolding predominantly stalled before reaching the 21 nm extension, highlighting the intrinsic defect of ΔF508 in achieving the fully folded state (Fig.3 J and K). Notably, the unfolding histograms had revealed lack of peaks within the initial 100 nm extension region for ΔF508 CFTR (Fig. 3C). As refolding occurs in reverse order of unfolding, this structural region corresponds to the 0–40 nm range in the refolding histogram (Fig. 2J, and 3 C and J). Supporting this interpretation, ΔF508 CFTR showed significantly reduced sampling below 40 nm compared to WT, underscoring the compromised stability of later-folding domains (Fig. 3J).

To dissect the folding defect of ΔF508 in details, we observed the occasional sampling of ΔF508 below 40 nm over extended periods and extracted transition rate constants to quantify which refolding steps were most affected by the ΔF508 (Fig. 3L). Contrary to expectations, the rate constant associated with NBD1 folding (81-to-63 nm transition), where F508 resides, remained largely unaffected (Fig. 3L). However, the forward transition from 33-to-21 nm, corresponding to the refolding of TM helices 11 and 12 in TMD2, was tenfold slower in ΔF508 CFTR (Fig. 3L, red arrow). Additionally, the backward rate, reflecting local unfolding, increased by twofold (SI Appendix, Fig. S10, right plots). The rate constant for the final transition, involving the refolding and assembly of NBD2 and lasso helices, could not be determined for ΔF508 due to the limited sampling beyond the 21 nm extension (Fig. 3L and SI Appendix, Fig. S10, right plots).

These observations were further substantiated by force-jump experiments (SI Appendix, Fig. S9 E–G). Analysis of the mapping broadness revealed local minima at 65, 80, and 105□nm (SI Appendix, Fig. S9F, bottom), which corresponded to 166, 198, and 260□nm in the 12-pN unfolding extension space (SI Appendix, Fig. S9G). These positions closely align with folding transitions associated with R domain, NBD1, and TMD1 respectively (Fig. 2J). The presence of refolding signatures corresponding to TMD1 and NBD1 suggests that these domains were able to complete tertiary structure formation, albeit with reduced stability. In contrast, folding transitions associated with TMD2 and NBD2 were significantly diminished or absent (Fig. 3C, J–L), indicating a defect in forming stable tertiary structures in the C-terminal half of the protein. Together, these results indicate that ΔF508 CFTR largely follows the same folding trajectory as WT CFTR but exhibits selective destabilization of intermediates beyond NBD1, particularly in the TMD2/NBD2 region.

Given that ΔF508 CFTR largely follows the same folding trajectory as WT CFTR, the impaired 33-to-21 nm transition suggests that the absence of F508 severely impairs the refolding of TM helix 11 and downstream structural elements (Fig. 2J and 3M). In the WT protein, the aromatic side chain of F508 interacts with hydrophobic residues in TM helix 11 (36). Our data suggest that these interactions are crucial for the proper folding of TM helix 11, and when TM helix 11 fails to fold correctly, the folding of downstream structural elements is also compromised (Fig. 3M) (17, 36, 53, 54). These observations strongly support a template-based folding pathway and suggest that the primary folding defect in ΔF508 lies in the inability of NBD1 to properly guide the folding of TM helix 11.

**Small-molecule correctors rescue the** Δ**F508 CFTR folding defect at 33-to-21 nm transition**

Having identified the impaired folding of TM helix 11 as the key defect in ΔF508 CFTR, we tested whether folding correctors could restore this transition. While ΔF508 in DMSO stalled beyond 40 nm during refolding, the addition of correctors enabled progression through the final transitions (Fig. 4A), ultimately achieving full refolding in a subset of molecules, more closely resembling WT behavior under the same treatment conditions (Fig. 4B). HMM analysis revealed that the intermediate states in corrector-treated ΔF508 overlapped substantially with those of WT, although residual instability at the 33-to-21 nm transition persisted under certain conditions (Fig. 4 C–F). Reflecting this, the overall refolding success probability remained lower than that of WT; nonetheless, the addition of ATP and a single corrector—or two correctors—significantly increased the probability of reaching the fully folded state (Fig. 4G). Notably, kinetic analysis revealed that lumacaftor and elexacaftor specifically accelerated the 33-to-21 nm transition, corresponding to the folding of TM helices 11 and 12, and restored the folding disrupted in ΔF508 (Fig. 4H, red arrow). These correctors also reduced the frequency of backward transitions, indicating a stabilization of folding intermediates (Fig. 4I, red arrow). Together, these findings confirm that the primary folding defect in ΔF508 lies in the inefficient refolding of TM helix 11 and its downstream elements, and demonstrate that folding correctors act by enhancing the folding rate at this critical step while preserving the folding pathway.

**Fig 4.**
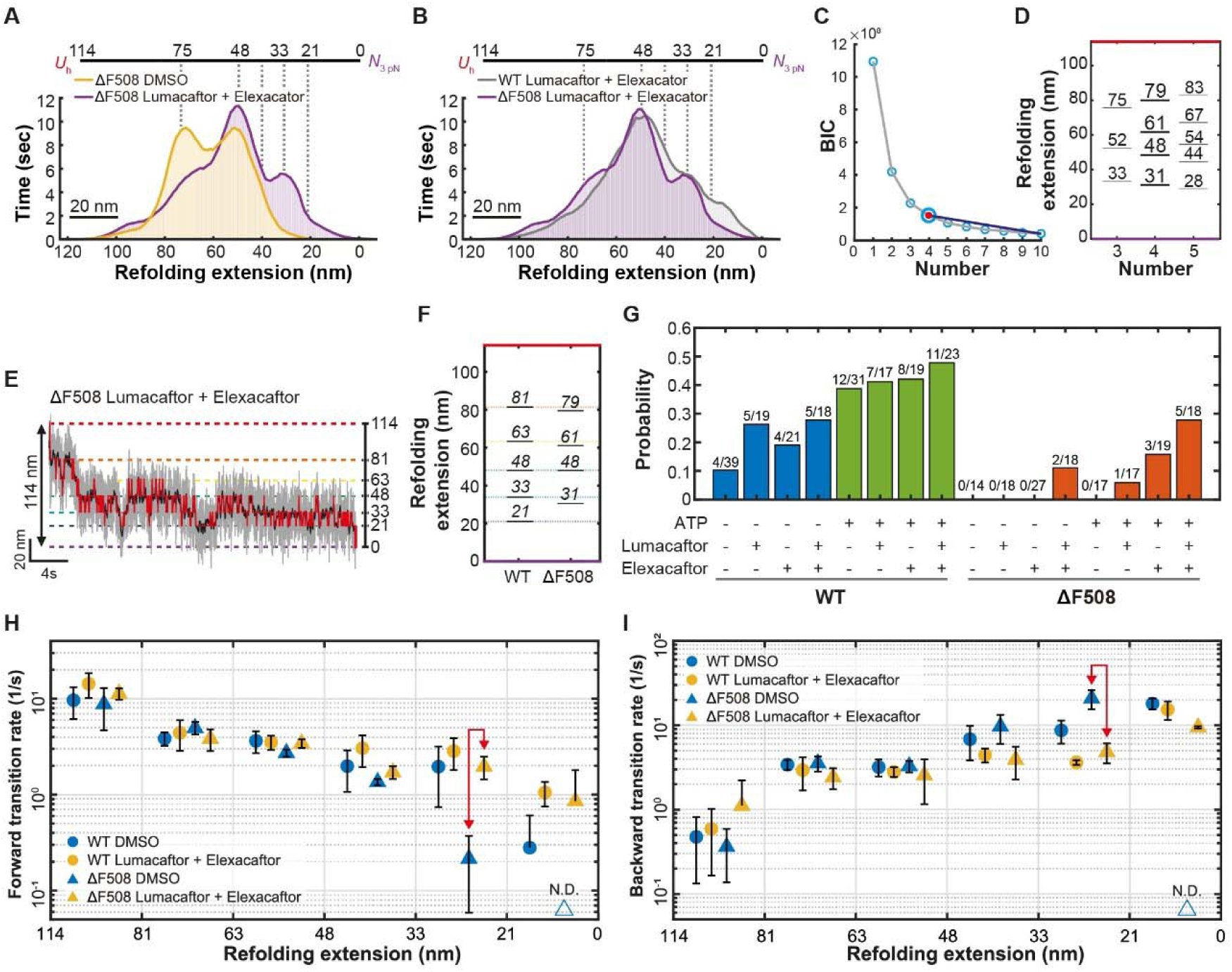
Folding impairment of ΔF508 CFTR and its correction by the folding correctors. (A) Refolding histograms at 3 pN for ΔF508 CFTR with or without folding correctors (*n* = 27 for ΔF508 DMSO, *n* = 39 for ΔF508 Lumacaftor + Elexacaftor). The refolding histogram for ΔF508 DMSO is replotted from Fig. 3J. (B) Refolding histograms at 3 pN for ΔF508 CFTR in comparison with WT CFTR, with both the folding correctors (*n* = 39 for ΔF508 Lumacaftor + Elexacaftor, *n* = 43 for WT Lumacaftor + Elexacaftor). The refolding histogram for ΔF508 Lumacaftor + Elexacaftor is replotted from Fig. 4A. (C and D) BIC analysis identifies four as the optimal number of intermediate states in ΔF508 CFTR refolding traces, one fewer than that detected for WT CFTR (C). The extensions of folding intermediates based on various assumed state numbers in the ΔF508 CFTR refolding traces (D). The analysis for ΔF508 CFTR was performed using all the conditions obtained using ΔF508 CFTR. (E) Representative refolding traces for ΔF508 CFTR with both lumacaftor and elexacaftor at 3 pN. 1.2 kHz raw data is gray, 10-Hz median filtered data is black, and the Viterbi path was overlaid on the data in red. Horizontal lines indicate extension values shown on the right. (F) HMM defined refolding intermediate states for WT and ΔF508 CFTR. The HMM defined refolding intermediate states for WT is replotted from Fig. 2C. (G) Refolding probabilities for WT and ΔF508 CFTR under the indicated conditions. The numbers above the bar graphs indicate the fraction of successfully refolded traces per total traces (successful over total trials). The first four data points are replotted from SI Appendix, Fig. S3A. (H) Forward transition rates for WT and ΔF508 CFTR, with and without lumacaftor and elexacaftor. Data represent means and standard deviations for *n* individual traces. *n* = 29 for WT Lumacaftor + Elexacaftor and *n* = 39 for ΔF508 Lumacaftor + Elexacaftor. The transition rates for WT DMSO and ΔF508 DMSO are replotted from Fig. 3L. N.D., not determined. (I) Backward transition rates for WT and ΔF508 CFTR under the same conditions. N.D., not determined.

We next undertook a quantitative analysis to delineate how the ΔF508 mutation and its pharmacological correction reshape the CFTR folding energy landscape. To this end, we determined 228 rate constants—encompassing both forward and backward transitions—across various conditions (SI Appendix, Fig. S10). These kinetic parameters enabled precise estimation of changes in free energy barriers (ΔΔ*G*^‡^) and relative shifts in the free energies of refolding intermediates (ΔΔ*G*_F_) (SI Appendix, Fig. S11). Conceptually, increased forward rates reflect catalytic enhancement of folding transitions, while decreased backward rates indicate stabilization of folded conformations (SI Appendix, Fig. S12A)—findings that were consistent with the dwell-time changes observed in the unfolding histogram analyses above. Therefore, we attributed variations in forward rates primarily to changes in ΔΔ*G*^‡^, and those in backward rates to a combination of altered ΔΔ*G*^‡^ and ΔΔ*G*_F_ (relative to WT CFTR in DMSO, unless otherwise specified) (Fig. 5A). Notably, because the free energy states of intermediate conformations are inherently linked across folding steps, ΔΔ*G*_F_ values derived from an earlier transition worked as initial conditions for calculating ΔΔ*G*^‡^ and ΔΔ*G*_F_ in the subsequent step (Fig. 5A and Methods for free energy analysis). This cumulative dependency underscored the necessity of determining rate constants for the full sequence of folding transitions, rather than interpreting each transition in isolation (SI Appendix, Fig. S11).

**Fig 5.**
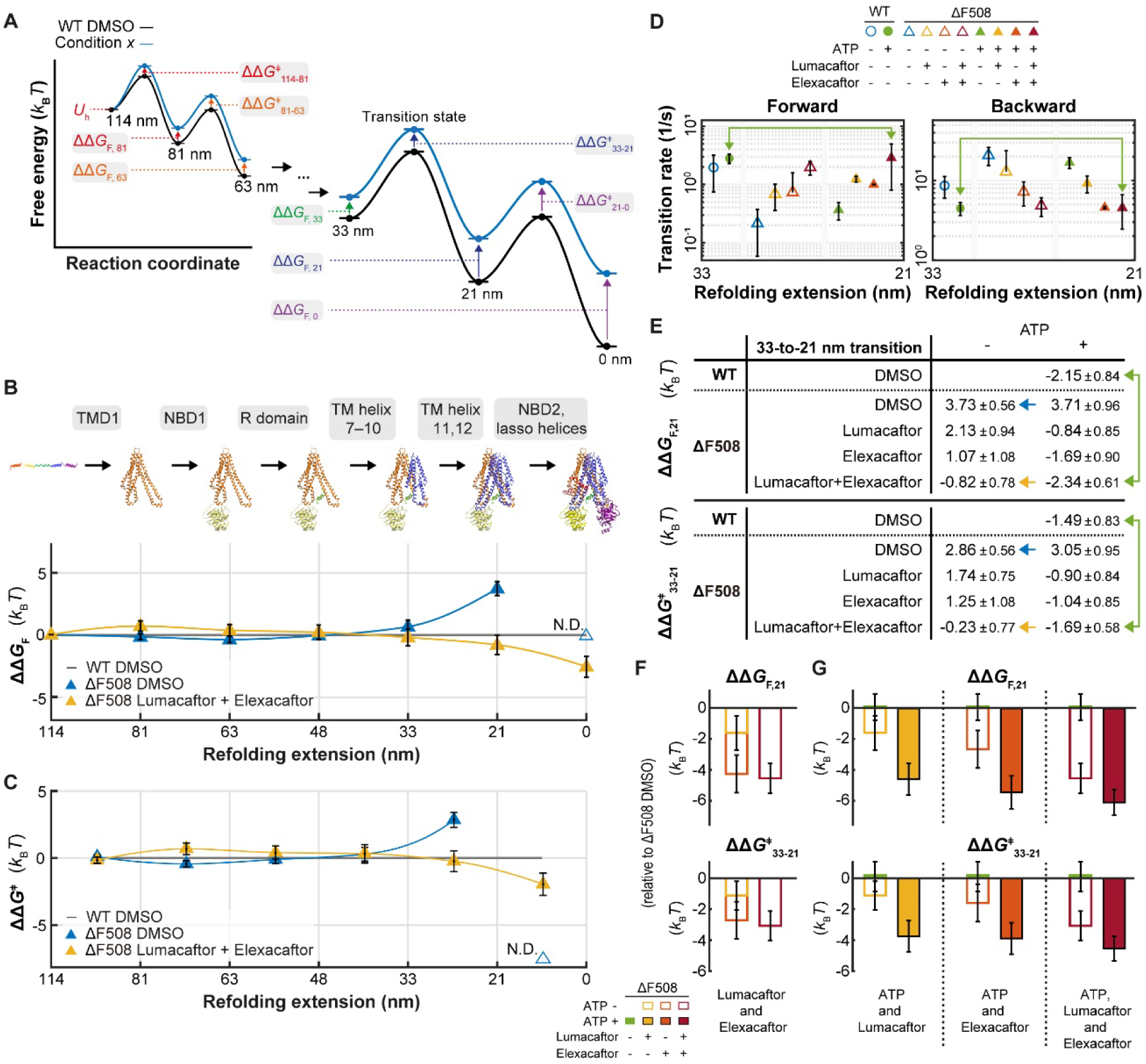
Small-molecule correctors rescue the ΔF508 CFTR folding defect at 33-to-21 nm transition. (A) Schematic representation of ΔΔ*G*^‡^ and ΔΔ*G*_F_, where the black curve represents the free energy landscape of WT CFTR in DMSO, and the blue curve represents the free energy landscape in the condition being examined. (B) ΔΔ*G*_F_ calculated relative to WT DMSO (gray line at ΔΔ*G*_F_ = 0), comparing ΔF508 DMSO and ΔF508 with Lumacaftor + Elexacaftor across all refolding intermediates. Data represent means and propagated errors calculated from transition rate values obtained from *n* individual traces (*n* values in SI Appendix, Fig. S10). N.D., not determined. (C) ΔΔ*G*^‡^ calculated relative to WT DMSO (gray line at ΔΔ*G*^‡^ = 0), comparing ΔF508 DMSO and ΔF508 with Lumacaftor + Elexacaftor across all refolding intermediates. Data represent means and propagated errors calculated from transition rate values obtained from *n* individual traces (*n* values in SI Appendix, Fig. S10). N.D., not determined. (D) Forward and backward transition rates for the 33-to-21 nm transition for WT and ΔF508 CFTR, with various combinations of lumacaftor, elexacaftor and ATP. Data represent means and standard deviations for *n* individual traces, with exact *n* values provided in SI Appendix, Fig. S10. (E) ΔΔ*G*_F,21_ and ΔΔ*G*^‡^_33-21_ values for WT and ΔF508 CFTR, calculated relative to WT DMSO. Data represent means and propagated errors for *n* individual traces, with exact *n* values provided in SI Appendix, Fig. S10. (F) Comparative analysis of ΔΔ*G* values at 33-to-21 nm transition to evaluate the combinatorial effects of lumacaftor and elexacaftor in rescuing the ΔF508 mutation. ΔΔ*G* values are calculated relative to ΔF508 DMSO. Error bars represent propagated errors. (G) Comparative analyses of ΔΔ*G* values at 33-to-21 nm transition to evaluate the combinatorial effects of ATP and the folding correctors in rescuing the ΔF508 mutation. ΔΔ*G* values are calculated relative to ΔF508 DMSO. Error bars represent propagated errors.

Using this approach, we systematically mapped how ΔΔ*G*_F_ and ΔΔ*G*^‡^ accumulated along the folding pathway for both WT and ΔF508 CFTR, and how these energy profiles were modulated by folding correctors and ATP (Fig. 5 B, C and SI Appendix, Fig. S11). The ΔF508 mutation severely destabilized the 21-nm refolding intermediate (ΔΔ*G*_F,21_), increasing its free energy by 3.73 *k*_B_*T* (Fig. 5 B, D, and E, blue symbols and an arrow). This destabilization propagated through the folding pathway in an amplified manner, ultimately rendering the free energy of the fully folded state (ΔΔ*G*_F,0_) undeterminable under the mutant condition (Fig. 5B). In addition, the ΔF508 mutation impaired the 33-to-21 nm transition itself, elevating the corresponding free energy barrier (ΔΔ*G*^‡^_33-21_) by 2.86 *k_B_T* (Fig. 5 C–E, blue symbols and an arrow). While either lumacaftor or elexacaftor alone partially mitigated these defects, their combined application effectively restoredΔΔ*G*_F,21_ and ΔΔ*G*^‡^_33-21_ to −0.82 *k_B_T* and −0.23 *k_B_T*, respectively, values closely aligning with those of WT CFTR in DMSO (Fig. 5 B–E, yellow symbols and arrows; WT reference shown in SI Appendix, Fig. S12 B and C).

Notably, in reducing ΔΔ*G*_F,21_ and ΔΔ*G*^‡^_33-21_, lumacaftor and elexacaftor exerted additive effects, with the effects being especially pronounced in ΔF508 CFTR (Fig. 5F; WT data shown in SI Appendix, Fig. S12D). This mechanistic synergy likely underlies the basis for the most widely used CFTR modulator therapy, which combines lumacaftor (or tezacaftor) and elexacaftor with the potentiator ivacaftor (55). Furthermore, lumacaftor, elexacaftor and their combination all exhibited strong synergy with ATP in promoting the 33-to-21 nm transition (Fig. 5G; WT data shown in SI Appendix, Fig. S12E). Structural studies suggest that lumacaftor binds to TMD1 (37), whereas elexacaftor binds the lasso helices and TM helices 10 and 11 (36). Thus, our data, combined with existing structural insights, suggest that lumacaftor, elexacaftor and ATP bind distinct structural elements to effect additive rather than redundant rescue of folding and interdomain assembly. The combined application of all three restored the forward and backward 33-to-21 nm transition rates of ΔF508 CFTR to values comparable to those determined for WT CFTR in the presence of ATP alone (Fig. 5 D and E, green arrows), achieving a near-complete rescue of this critical transition.

### Rescue of the terminal folding step in WT and ΔF508 CFTR

We next examined the final 21-to-0 nm transition, which corresponds to the folding of NBD2 and the lasso helices. Lumacaftor, elexacaftor, ATP and their combinations exhibited pronounced restorative effects on this transition in both WT and ΔF508 CFTR (Fig. 6 A–C). These findings are consistent with our earlier observation that this step represents the slowest and most recalcitrant transition, even in WT CFTR (Fig. 2K), and therefore possesses the greatest potential for energetic improvement (Fig. 6 D, E and SI Appendix, Fig. S11).

**Fig 6.**
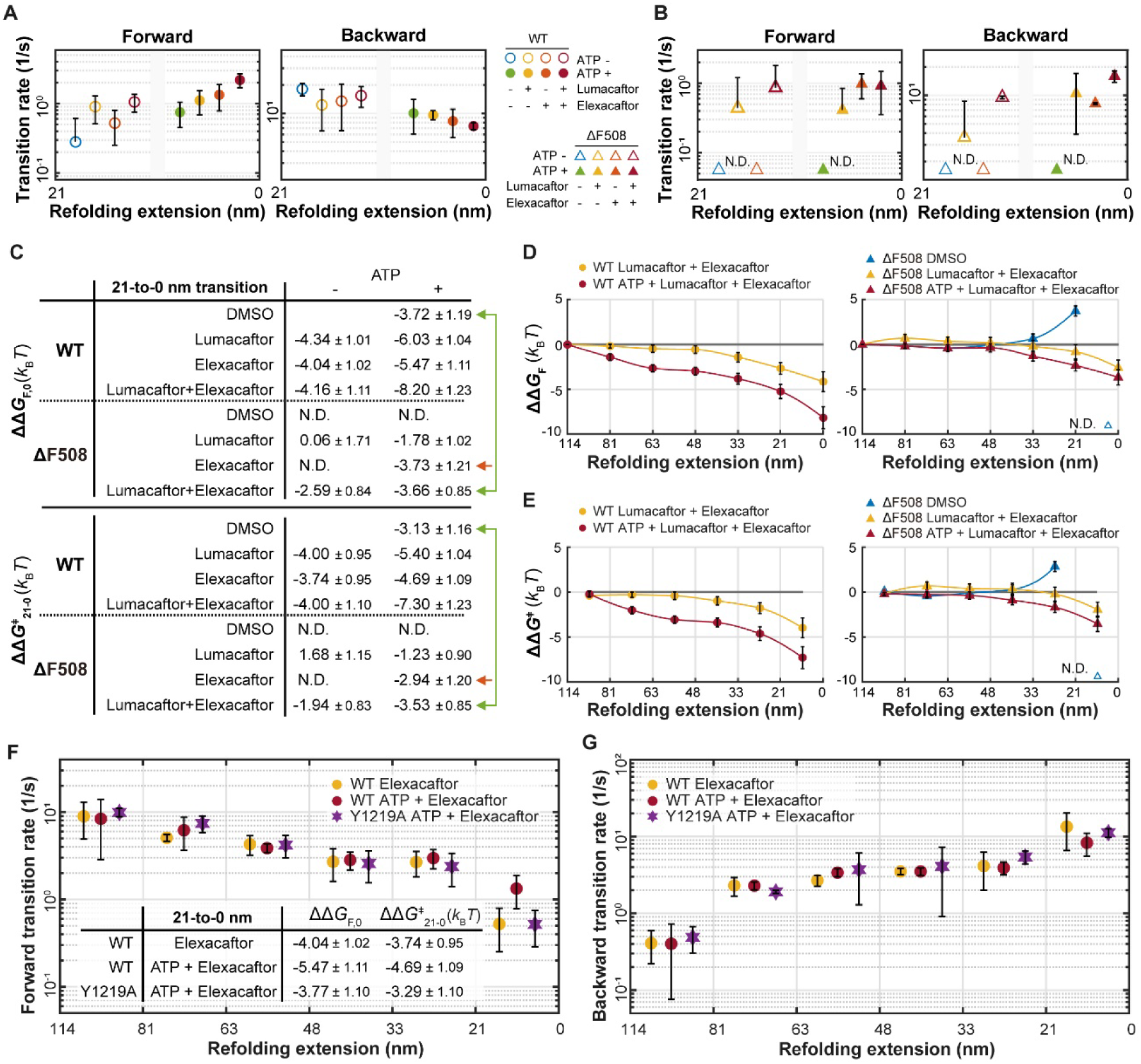
Small-molecule correctors rescue the ΔF508 CFTR folding defect at 33-to-21 nm transition. (A) Forward and backward transition rates for the 21-to-0 nm transition for WT with various combinations of lumacaftor, elexacaftor and ATP. Data represent means and standard deviations for *n* individual traces, with exact *n* values provided in SI Appendix, Fig. S10. (B) Forward and backward transition rates for the 21-to-0 nm transition for ΔF508 with various combinations of lumacaftor, elexacaftor and ATP. Data represent means and standard deviations for *n* individual traces, with exact *n* values provided in SI Appendix, Fig. S10. (C) ΔΔ*G*_F,0_ and ΔΔ*G*^‡^_21-0_ values for WT and ΔF508 CFTR, calculated relative to WT DMSO. Data represent means and propagated errors for *n* individual traces, with exact *n* values provided in SI Appendix, Fig. S10. (D) ΔΔ*G*_F_ calculated relative to WT DMSO (gray line at ΔΔ*G*_F_ = 0), comparing WT Lumacaftor + Elexacaftor and WT ATP + Lumacaftor + Elexacaftor (left), ΔF508 Lumacaftor + Elexacaftor and ΔF508 ATP + Lumacaftor + Elexacaftor (right), across all refolding intermediates. Data represent means and propagated errors calculated from transition rate values obtained from *n* individual traces (*n* values in SI Appendix, Fig. S10). (E) ΔΔ*G*^‡^ calculated relative to WT DMSO (gray line at ΔΔ*G*^‡^ = 0), comparing WT Lumacaftor + Elexacaftor and WT ATP + Lumacaftor + Elexacaftor (left), ΔF508 Lumacaftor + Elexacaftor and ΔF508 ATP + Lumacaftor + Elexacaftor (right), across all refolding intermediates. Data represent means and propagated errors calculated from transition rate values obtained from *n* individual traces (*n* values in SI Appendix, Fig. S10). (F and G) Forward (F) and backward (G) transition rates for refolding transitions of Y1219A and WT CFTR with the indicated conditions. Data represent means and standard deviations for *n* individual traces. *n* = 45 for Y1219A ATP + Elexacaftor. The transition rates for WT Elexacaftor and WT ATP + Elexacaftor are replotted from SI Appendix, Fig. S10. ΔΔ*G*_F,0_ and ΔΔ*G*^‡^_21-0_ values for Y1219A, compared with WT CFTR (F, inset).

As the destabilizing effect of the ΔF508 mutation propagated through the folding pathway, the stabilizing influence of folding correctors on refolding intermediates also accumulated progressively in WT CFTR (Fig. 6 D, E and SI Appendix, Fig. S11A). In the absence of ATP, the effects of the correctors became evident beyond the 48 nm extension (Fig. 6 D, E and SI Appendix, Fig. S11A, yellow circles), corresponding to TMD2 and subsequent domains. ATP further amplified these stabilizing effects, culminating in the largest ΔΔ*G* reduction at the 21-to-0 nm transition and highlighting the synergistic interplay between correctors and ATP (Fig. 6 D, E and SI Appendix, Fig. S11A, red circles). Notably, in ΔF508 CFTR, the cumulative energetic gains from folding correctors and ATP were quantitatively smaller than those observed in WT CFTR (Fig. 6 D and E, compare left and right panels). This underscores the importance of the structural integrity of pre-formed domains in effectively propagating the effects of correctors through the folding pathway. Collectively, these observations support a template-driven model in which the effects of folding correctors propagate through downstream folding transitions, rather than acting solely at isolated structural elements.

This model was further supported by detailed energetic analysis of the 21-to-0 nm transition. The large negative values of ΔΔ*G*_F_,_0_ and ΔΔ*G*^‡^_21-0_ determined for WT CFTR indicated that correctors and ATP actively catalyzed this final transition, rather than merely accelerating preceding steps (Fig. 6C). Although baseline ΔΔ*G* values could not be measured for ΔF508 CFTR in the presence of DMSO due to the paucity of complete refolding events, the energetic effects of pharmacological correctors could still be inferred. Notably, while elexacaftor alone had minimal effect on the 21-to-0 nm transition, its combination with ATP reduced ΔΔ*G*_F_,_0_ by −3.73 *k*_B_*T* and ΔΔ*G*^‡^_21-0_ by −2.94 *k_B_T* (Fig. 6C, orange arrows). Importantly, this synergy was abolished by the Y1219A substitution in WT CFTR, which disrupts ATP binding to NBD2 (Fig. 6 F and G), indicating that elexacaftor’s effect depends on direct ATP interaction with NBD2 rather than an allosteric influence from NBD1. Although elexacaftor binds distally from NBD2, these results underscore the critical role of interdomain communication in CFTR folding and reveal that effective pharmacological rescue requires not only local stabilization but also long-range energetic coupling between structural domains.

Finally, in the presence of both folding correctors and ATP, the ΔΔ*G*_F_,_0_ and ΔΔ*G*^‡^_21-0_ values for ΔF508 CFTR reached −3.66 and −3.53 *k*_B_*T*, respectively, bringing these values close to those observed for WT CFTR with ATP (Fig. 6C, green arrows). While the synergistic effects of lumacaftor and elexacaftor, in the presence of ATP, appeared attenuated during the folding of NBD2 and the lasso helices (Fig. 6C, compare orange and green arrow; WT data shown in SI Appendix, Fig. S12 F and G)—possibly reflecting a convergence of their mechanisms at this terminal stage—the final and rate-limiting folding transition was fully restored to WT-like levels through the combined action of folding correctors and ATP.

## Discussion

In this study, we examined the unfolding and refolding processes of individual CFTR molecules in lipid bicelles. We began with a fully folded CFTR molecule, applied mechanical force to completely unfold it, and then relaxed the force to observe its refolding. This process differs substantially from *in vivo* CFTR biosynthesis, where folding occurs through both co-translational and post-translational phases and involves a network of molecular chaperones and protein quality control mechanisms (30, 56–58). Nevertheless, we show that native-like CFTR structures can be achieved in *in vitro* refolding experiments and the probability of successful refolding is sensitive to ATP and folding correctors. These data provide direct support for Anfinsen’s thermodynamic hypothesis that protein structure is determined solely by its amino acid sequence (59). This finding is particularly significant given CFTR’s large, multidomain architecture.

By quantitatively analyzing thermodynamic and kinetic aspects of the CFTR refolding pathway, we propose that CFTR exhibits a hierarchical, templated folding pathway where the N-terminal TMD1 and NBD1 fold first and act as scaffolds for the subsequent folding of their C-terminal counterparts, TMD2 and NBD2. Consequently, TMD2 and NBD2 integrate directly into the growing CFTR structure, without a distinct final assembly phase between the N- and C-terminal domains. The rate-limiting step of the entire process is the folding of NBD2 and the N-terminal lasso helices. Although ΔF508 CFTR follows the same pathway as WT CFTR, the lack of F508 at the NBD1/TMD2 interface reduces the stability of NBD1, but more importantly, it slows the folding of TMD2 by an order of magnitude. Furthermore, weakened NBD1/TMD2 interface in ΔF508 CFTR destabilizes both TMD2 and NBD2, even after folding is complete, as evidenced by the total disappearance of early unfolding peaks within the first 100 nm extension range at 12 pN. These defects ultimately obstruct the formation of the native conformation and weaken structural connectivity across CFTR.

Although our proposed template-driven model deviates from the traditional view that individual CFTR domains fold separately before assembling into the native-like conformation (30, 58), it aligns well with many observations reported in the literature. During CFTR biosynthesis, as the polypeptide chain emerges from ribosome, individual domains—except for NBD2—rapidly adopt conformations that render them resistant to proteolysis (16, 53, 58, 60). By contrast, NBD2 folding is not only slow but also dependent on the presence of other domains (30, 61–63). Furthermore, in the early co-translational stage, NBD1 already interacts with TMD1 (64, 65) and the formation of the most stable TMD2 conformation requires both TMD1 and NBD1 (30). These findings support that in the cellular environment, CFTR folding also follows a template-driven pathway rather than an independent domain-folding model.

Crucially, this templated folding mechanism implies that the free energy states of individual folding intermediates are intimately coupled along the CFTR folding trajectory. Leveraging our single-molecule MT assay, we determined 228 forward and backward transition rates spanning distinct folding steps under various pharmacological and ATP conditions. Although all kinetic measurements were necessarily performed under finite mechanical load, these transition rates were measured at the lowest force compatible with reliable resolution, which likely brackets the mechanical regime experienced by nascent membrane proteins during translation and chaperone-assisted folding. This comprehensive dataset enabled precise delineation of relative free energy shifts across the entire CFTR folding energy landscape. These measurements provided an energetic perspective on how the ΔF508 mutation and folding correctors modulate specific folding transitions—insights difficult to capture through bulk biochemical or cellular assays, which predominantly reflect terminal folding outcomes. The resulting folding energy landscape revealed that the impact of the ΔF508 mutation was not confined to NBD1 but instead propagated throughout the entire folding pathway. Notably, the destabilizing effect became increasingly amplified beyond 33□nm, affecting transitions associated with TMD2, NBD2, and the lasso helices. This finding underscores a cascading mechanism whereby an initially localized defect exerts widespread energetic disruptions across the multidomain architecture of CFTR. In parallel, the effects of folding correctors were also transduced along the folding trajectory and were further potentiated by ATP. Somewhat unexpectedly, the cumulative energetic gains from folding correctors and ATP in ΔF508 CFTR were markedly smaller than those observed for WT CFTR, emphasizing the critical role of structural integrity in early domains for the effective propagation of pharmacological rescue. Nevertheless, the combined action of lumacaftor and elexacaftor—particularly when administered with ATP—robustly enhanced the final folding transitions in ΔF508 CFTR, ultimately restoring their energetics to levels comparable to WT.

Our data further reveal that folding correctors act not only by stabilizing folded domains but also by actively catalyzing specific folding transitions, as evidenced by reductions in the energy barriers. This dual mode of action suggests that folding correctors and ATP serve as allosteric modulators that reshape the folding energy landscape—functioning in a chaperone-like manner to facilitate otherwise inefficient transitions, rather than merely locking pre-existing folded conformations. Furthermore, the quantitative synergy analysis suggests that lumacaftor, elexacaftor, and ATP engage distinct structural elements and likely reinforce separate interfaces critical for the assembly of its C-terminal counterparts. Interestingly, the synergistic action between lumacaftor and elexacaftor was diminished during the final folding step involving NBD2 and the lasso helices, possibly reflecting a convergence of their actions at this terminal stage. Together, these findings support a multi-faceted mechanism, wherein distinct yet complementary modulators can act in concert with ATP to restore native folding—highlighting a therapeutic strategy rooted in energetic complementation.

The experimental platform described here offers a powerful framework for interrogating how rare CFTR mutations—particularly those unresponsive to current correctors and potentiators—disrupt the folding process, pinpointing the folding transitions that a given set of mutations imposes its most severe energetic penalty. As demonstrated, the system is amenable to testing novel corrector candidates and quantifying their ability to resuscitate specific folding transitions. When combined with established cell-based and in vivo models, these insights can inform the mechanistic understanding of how specific mutations perturb folding energetics and how correctors restore function—thereby laying the groundwork for extending CFTR-targeted therapies and advancing treatments for other protein folding disorders.

## Materials and Methods

### Expression and purification of the human CFTR

Dual-SpyTagged CFTR fused to a C-terminal PreScission Protease (PPX)-cleavable GFP tag was cloned into the BacMam vector and expressed as previously described for the WT CFTR (6). P3 baculovirus was generated as previously described (66) using Sf9 cells (Gibco, catalogue number 11496015, lot number 1670337) cultured in sf-900 SFM medium (Gibco), supplemented with 5% (v/v) heat-inactivated fetal bovine serum (FBS) and 1% (v/v) antibiotic–antimycotic (Gibco).

HEK293S GnTI^−^ (ATCC CRL-3022, lot number 62430067) cells were cultured in suspension using FreeStyle 293 medium (Gibco) supplemented with 2% (v/v) heat-inactivated FBS and 1% (v/v) antibiotic–antimycotic (Gibco), shaking at 37□°C with 8% CO_2_ and 80% humidity. At a density of 2.8□×□10^6^□cells/mL, cells were infected with baculovirus. 10 mM sodium butyrate was added to the culture after 12 hours and the incubation temperature was reduced to 30□°C. After 48 hours, the cells were pelleted and flash-frozen in liquid nitrogen.

For expression of ΔF508 CFTR, the culture medium was supplemented with 100 nM lumacaftor and 30 nM elexacaftor, and the incubation temperature after sodium butyrate addition was adjusted to 32 °C.

For protein purification, cells were solubilized in extraction buffer containing 1.25% (w/v) lauryl maltose neopentyl glycol (LMNG), 0.25% (w/v) cholesteryl hemisuccinate (CHS), 200□mM NaCl, 20□mM HEPES (pH□7.2 with NaOH), 2□mM MgCl_2_, 2 mM dithiothreitol (DTT), 20% (v/v) glycerol, 1□μg/mL pepstatin A, 1□μg/mL leupeptin, 1□μg/mL aprotinin, 100□μg/mL soy trypsin inhibitor, 1□mM benzamidine, 1□mM phenylmethylsulfonyl fluoride (PMSF) and 3□µg/mL DNase I for 75□min at 4□°C. The detergent extract was clarified by centrifugation at 75,000g for 2□×□20 min at 4□°C. The clarified lysate was mixed with NHS-activated Sepharose 4 Fast Flow resin (GE Healthcare) that had been conjugated with a GFP nanobody. After 1 hour, the resin was packed into a gravity flow chromatography column and then washed with 20 column volumes of wash buffer containing 0.06% (w/v) digitonin, 200□mM NaCl, 20□mM HEPES (pH 7.2 with NaOH), 2 mM DTT, and 2□mM MgCl_2_. The resin was then incubated for 2□h at 4□°C with 0.35□mg/mL PPX to cleave off the GFP tag. The eluate was collected by dripping through Glutathione Sepharose 4B resin (Cytiva), concentrated and loaded onto a Superose 6 10/300 GL column (GE Healthcare), equilibrated with 200 mM NaCl, 20 mM HEPES (pH 7.2 with NaOH), 2 mM MgCl_2_, 2 mM DTT, and 0.06 % (w/v) digitonin. Peak fractions were pooled, concentrated to 2 µM, aliquoted, and flash-frozen in in liquid nitrogen.

For purification of the Y109C/E1126C CFTR variant, DTT was omitted from all buffers. For purification of the ΔF508 CFTR variant, 1 mM ATP was added to extraction and wash buffers. A dialysis step was added prior to gel filtration to remove lumacaftor and elexacaftor copurified with CFTR from the culture medium. The eluted sample was dialyzed for a total of 48 hours against wash buffer with several buffer exchanges.

### Patch-Clamp Electrophysiology

Chinese hamster ovary (CHO-K1) cells (ATCC CCL-61, lot number 70014310) were maintained in DMEM-F12 (ATCC) supplemented with 10% (v/v) heat-inactivated FBS and 1% (v/v) GlutaMAX (Gibco) at 37□°C. The cells were plated in 35-mm cell culture dishes (Falcon) 24□hours before transfection. Transfection with dual-SpyTagged and C-terminally GFP-fused CFTR was performed, using Lipofectamine 3000 according to the manufacturer’s protocol (Invitrogen). Culture medium was replaced with DMEM-F12 supplemented with 2% (v/v) heat-inactivated FBS and 1% (v/v) GlutaMAX 8□hours following transfection. The cells were incubated at 30□°C for 24□hours before recording.

Recordings were performed using the inside-out patch configuration with local perfusion at the patch. Bath solution was 145□mM NaCl, 2□mM MgCl_2_, 5□mM KCl, 1□mM CaCl_2_, 5□mM glucose, 5□mM HEPES and 20□mM sucrose (pH 7.4 with NaOH). Pipette solution was 140□mM NMDG, 5□mM CaCl_2_, 2□mM MgCl_2_ and 10□mM HEPES (pH 7.4 with HCl). Perfusion solution was 150□mM NMDG, 2□mM MgCl_2_, 1□mM CaCl_2_, 10□mM EGTA and 8□mM Tris (pH 7.4 with HCl). Dual-SpyTagged CFTR was activated by PKA (Sigma-Aldrich) phosphorylation with 3□mM ATP.

Borosilicate glass pipettes were pulled from capillaries (outer diameter 1.5□mm, inner diameter 0.86□mm, Sutter) to 1.5–2.5 MΩ resistance and fire polished. Membrane potential was clamped at −30□mV. Macroscopic currents were recorded at 25□°C using an Axopatch 200B amplifier, a Digidata 1550 digitizer and the pClamp software suite (Molecular Devices). Recordings were low-pass-filtered at 1□kHz and digitized at 20□kHz. The displayed recordings were further low-pass filtered at 100□Hz. Data were analyzed with Clampfit.

### Disulfide-dependent gel shift analysis

24 hours before transfection, HEK293S GnTI^−^ cells were seeded in Corning CellBIND 6-well plates in DMEM-F12 medium (ATCC) supplemented with 10% (v/v) heat-inactivated FBS and 1% (v/v) GlutaMAX (Gibco) at 37□°C. Transfection with C-terminally GFP-fused cysteine variants of CFTR was performed using Lipofectamine 3000 according to the manufacturer’s protocol (Invitrogen). At 8 hours after transfection, culture medium was replaced with DMEM-F12 supplemented with 2% (v/v) heat-inactivated FBS and 1% (v/v) GlutaMAX. The cells were incubated at 30□°C for an additional 24□hours, harvested by resuspension in Dulbecco’s phosphate-buffered saline (Gibco), pelleted by centrifugation, and snap-frozen in liquid nitrogen. Cells were lysed by resuspension in extraction buffer containing 1.25% (w/v) LMNG, 0.25% (w/v) CHS, 200 mM NaCl, 20 mM HEPES (pH 7.2 with NaOH), 2 mM MgCl_2_, 20% (v/v) glycerol, 1 μg/mL pepstatin A, 1 μg/mL leupeptin, 1 μg/mL aprotinin, 100 μg/mL soy trypsin inhibitor, 1 mM benzamidine, 1 mM PMSF and 3 µg/mL DNase I, and incubated for 1 hour at 4 °C. Lysates were clarified, separated by SDS-PAGE, and the gel imaged for GFP fluorescence.

### Preparation of DNA handles

The preparation of SpyCatcher-conjugated DNA followed an established protocol (42, 44). In summary, 512-base-pair DNA fragments carrying biotin or digoxigenin modification at one end and an amine group at the other were amplified by PCR. These DNA fragments were reacted with SM(PEG)_2_ (a PEGylated SMCC crosslinker; ThermoFisher Scientific) using an amine-sulfhydryl crosslinking reaction for 1 hour at room temperature. Following DNA maxiprep purification, the fragments were conjugated to purified SpyCatcher/Maltose Binding Protein (MBP) through a thiol-maleimide crosslinking reaction, which proceeded overnight at 4°C. To eliminate unconjugated SpyCatcher and unconjugated DNA, the resultants were purified using anion exchange chromatography with a 1 mL Mono Q column (GE Healthcare) followed by amylose affinity chromatography (New England BioLabs). The final SpyCatcher-DNA handles, stored in 50 mM Tris (pH 7.5) and 150 mM NaCl, were concentrated to approximately 200 nM and preserved in 5 µl aliquots at −80°C.

### Instrumentation of single-molecule MT

A custom-built MT system assembled on an inverted microscope (Olympus Live Cell Instrument) was used as previously described (41). A pair of permanent neodymium magnets was positioned using a translational stage (Physik Instrumente) to apply forces ranging from ∼10 fN to 50 pN. A super-luminescent diode (λ = 680 nm, Qphotonics) provided illumination, generating diffraction patterns for both magnetic and reference beads. These images were captured at rates of up to 1.2 kHz using a high-speed CMOS camera (Mikrotron). Calibration tables for individual beads were created by recording diffraction patterns while adjusting the objective lens position with a piezoelectric nano-positioner (Mad City Labs). The 3D positions of magnetic beads were tracked in real time by comparing diffraction patterns with the pre-recorded calibration data. Single-molecule experiments were controlled using custom LabView software.

### Single-molecule MT experiments

WT or mutant CFTR proteins solubilized in 0.06% (w/v) digitonin were mixed with the SpyCatcher-DNA handles in 1.3% (w/v) bicelle buffer. The bicelles consisted of 70:30 ratio of DMPC and DMPG lipids (Avanti Polar Lipids) and CHAPSO (Sigma-Aldrich) at a 2.8:1 molar ratio in 50 mM Tris pH 7.5 and 150 mM NaCl buffer with 2 mM MgCl_2_ and 2 mM Tris (2-carboxyethyl) phosphine hydrochloride (TCEP). The mixture was incubated for 90 minutes at room temperature to attach DNA handles to both ends of the CFTR proteins, using a 2:1 molar ratio of CFTR protein to SpyCatcher-DNA handles. After incubation, the protein-DNA hybrid complexes were diluted to a final DNA handle concentration of approximately 5 nM. CFTR protein conjugated with two DNA handles was always freshly prepared before the MT experiment.

For the experiments, 0.02% (w/v) streptavidin-coated polystyrene particles (3.11 µm, Spherotech) were first injected into a homemade flow cell consisting of two cover slips (VWR No. 1.5). The bottom cover slip was coated with mPEG and biotin-PEG at a 100:3 molar ratio. After 7 minutes of incubation, unbound reference beads were removed by extensive microfluidic buffer exchange. Next, 15 nM Neutravidin was injected and incubated for 8 minutes. Following the removal of unbound Neutravidin, the chamber was washed with bicelle buffer prior to sample injection. The bicelle buffer was supplemented with ligands according to experimental conditions (e.g., ATP diluted in bicelle buffer for ATP conditions or lumacaftor diluted in bicelle buffer for lumacaftor conditions). The final ligand concentrations were: ATP, 1 mM; MgCl_2_, 2 mM; lumacaftor, 1 µM; elexacaftor, 1 µM. CFTR conjugated with DNA handles was then introduced into the chamber and incubated for 20 minutes, followed by buffer exchange to remove unbound protein. Additional washing steps were performed under specific conditions: in the ‘phosphorylated, ATP’ condition, an extra wash step was included to ensure complete removal of PKA, while in experiments using ΔF508 samples, additional washing was performed to eliminate any residual folding correctors from the purification step. Finally, magnetic beads (2.8 µm diameter, Invitrogen) were injected and incubated for 1 hour.

### Extensible Worm-Like Chain (eWLC) model for DNA and unstructured polypeptide

The FECs for DNA and unstructured polypeptides were analyzed using the eWLC model, which characterizes the mechanical response of semi-flexible biopolymers under applied tension. The eWLC equation is given by:

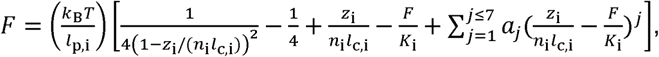

where the index *i* denotes either DNA or unstructured polypeptide (*p*). Here, *k*_B_*T* represents the thermal energy, *l*_c_ and *l*_p_ correspond to the contour length and persistence length, respectively (*l*_c,DNA_ = 0.338 nm, *l*_c,p_ = 0.36 nm and *l*_p,DNA_ = 38.5 nm, *l*_p,p_ = 0.39 nm). The elastic modulus values are *K*_p_∼50 µN and *K*_DNA_∼500 pN. *F* is the applied force and *a_j_* are polynomial coefficients for improved approximation. The total number of monomers in each component is defined as follows: *n*_DNA_ = 512 for each handle, *n*_linker,p_ = 54 for polypeptide linker between CFTR and the for improved approximation. The total number of monomers in each component is defined as DNA handle, and *n*_CFTR,p_ = 1426 for the CFTR protein.

### Kessler-Rabin (KR) model for rigid-like biopolymers

To describe the helical states, *U*_h_, the KR model was used,

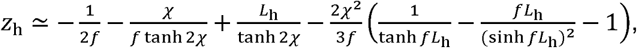

where 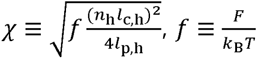 and *n_h_* is the number of amino acids consisting of the transmembrane helix. The persistence length (*l*_p,h_) is 9.17 nm and the contour length (*l*_c,h_) along helical axis is averagely 0.16 nm per amino acid.

### Extensions calculation in force-ramp and force-jump experiments

In force-ramp and force-jump experiments, the observed extension values can be estimated from a linear superposition of extensions of all components in tweezing system. The extensions expected for the *U*_c_ and *U*_h_ states are thus described as follows.

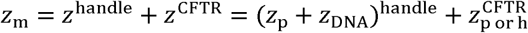

where *z*_m_ is the measured extension, *z*_p_ represents the extension of the unstructured polypeptide linker between DNA handle and the target protein (linkers from each end of the protein to SpyCatcher), *z*_DNA_ corresponds to the extension of the DNA in the DNA handle, and 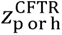 is the total molecular extension of CFTR with contributions from unstructured and/or helical parts. The *z*_p_ and *z*_DNA_ values are inversely calculated from the eWLC model at given force levels, and 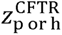 corresponds to the total molecular extension of CFTR. The values of 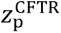 and 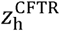 are derived from the eWLC and KR model, respectively. These calculations also incorporate the end-to-end distance of structured domains that contribute to the protein’s tertiary conformation. For instance, when stretching CFTR in its native state, 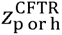 is replaced by a *d_N_* value of 6.5 nm, which is the end-to-end distance derived from the native state structure (PDB: 5UAK) (32).

To understand the relative changes in extension during high-force unfolding (i.e. to understand how the extension values correspond to which intermediate states), we calculated the number of amino acids that undergo stretching and the change in the end-to-end distance of the remaining structured regions. The increase in extension for an intermediate state 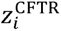 is proportional to the number of unfolded amino acids (Δ*n_i_*). This value is obtained by multiplying the number of unfolded amino acids by the extension per amino acid at a given force. In summary, the relationship used for calculating the measured extension value at an intermediate state *i*(*z*_m,i_) is:

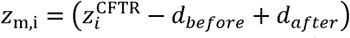

where *d_before_* and *d_after_* represent the end-to-end distance of the structured regions before and after the transition, respectively.

### Cross-correlation analysis

To assess the common effect of the ligands under different conditions (i.e. Lumacaftor effect in WT Lumacaftor and WT ATP + Lumacaftor), we applied a cross-correlation method by computing the product of corresponding peak values at each extension. A matrix was generated, where each element was calculated as:

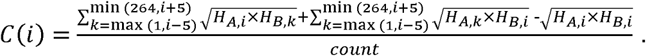

*C*(*i*) represents the correlation product at position *i* (0 < *i* < 264) and *H_X,i_* corresponds to the histogram height at position *i* for condition of *X* (*A* or *B*). The *count* is the number of elements included in the sum for normalization purposes. We included values within *i* ±5 nm to account for peak widths. Higher correlation products were expected when both histogram values were elevated. This observation indicates that, despite fluctuations inherent in high-force unfolding experiments, peaks consistently appeared in both histograms. We posited that these peaks represent the points at which structural domains, stabilized by a reagent common to both conditions, exhibit greater resistance to mechanical force.

### Sliding window difference analysis

To identify regions exhibiting ligand-induced changes in dwell time, we performed pairwise histogram subtraction between conditions differing by a single ligand (e.g., to assess the effect of Lumacaftor, the WT Lumacaftor histogram was subtracted from the WT□DMSO histogram). While ligand binding can have allosteric effects across the protein, its most direct and localized impact typically occurs near the binding site, often stabilizing specific TM helices. Lumacaftor is known to bind TM helices 1, 2, 3, and 6, and is therefore expected to most immediately influence the dwell time associated with TM helix 6. Since TM segments generally unfold as helical hairpins, and the typical extension of a hairpin at 12□pN is approximately 20□nm, we applied a 20-nm sliding window to the histograms to detect such localized effects.

### Area under the curve (AUC) comparison analysis

To statistically evaluate the impact of specific substances on CFTR unfolding dynamics, we employed an AUC analysis. For each condition, we randomly selected 40 unfolding cycles to generate a histogram, which was sufficient to observe divergence in the resulting histograms. This sampling process was repeated 100 times to calculate AUCs over defined extension ranges corresponding to distinct structural domains: 30–50 nm (NBD2 and lasso helices), 157–177 nm (NBD1), and 207–227 nm (TMD1). The boxplot elements represent the following: median (red vertical line); box limits (25^th^ and 75^th^ percentiles); whiskers (extending to 1.5×IQR). Statistical significance was tested using a two-sample t-test (***p ≤ 1 × 10^−3^).

### Hidden Markov Model analysis

HMM analysis was employed to determine the folding/unfolding intermediate states from the time-resolved low-force extension traces recorded at 1.2 kHz. The adjustable parameters in our system are the number of states (*n*), the extension position for *i*-th intermediate state (*z*□*i*;), and the transition matrix of rates between states 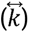. The optimal number of states (*n*) was obtained from BIC: BIC=*q*ln(*N*)−2ln(*L*□) where *q* is the number of output parameters given by model, *N* is sample size and *L*□ is the maximum value of the likelihood function. Maximum likelihood estimation was performed using the Baum-Welch algorithm. BIC as a function of the number of states determines the optimal number by finding the point where the BIC slope substantially changes. The extension traces were median-filtered with 10-Hz window, and the extension position/deviation for each state was estimated from the Gaussian Mixture Model (GMM) in the HMM analysis. The rates (*i.e.*, the transition matrix) were then determined using the optimal parameters for the number of states and extension positions. Finally, the resulting traces were verified by the Viterbi algorithm.

### Force jump analysis

To investigate the correlation between intermediates observed at 12 pN and 3 pN, force-jump experiments were performed by rapidly increasing and decreasing the force between 3 pN and 12 pN while monitoring the magnetic bead’s position in the extension space. To ensure precise bead positioning, extensions were recorded at each force, and histograms were generated with Gaussian fitting. The peak of each histogram was used as the representative extension value for comparisons between 3 pN and 12 pN. To quantify the correlation between extension values at these forces, a 20-nm moving window was applied across the 3-pN extension space. This window size was chosen based on the minimum extension observed for an individual domain, which was approximately 20 nm. A box plot was constructed using the 25^th^ to 75^th^ percentile range, and IQR values were analyzed to assess variability.

### Free energy analysis

To assess the impact of forward and backward transition rates on the free energy changes of intermediate and transition states, relative free energy differences (ΔΔ*G*^‡^ and ΔΔ*G*_F_) were calculated with respect to WT CFTR under DMSO conditions. Free energy differences for each transition from the *n*^th^ state, *i_n_* nm to the (*n*+1)^th^ state, *i*_n+1_ nm were determined iteratively using:

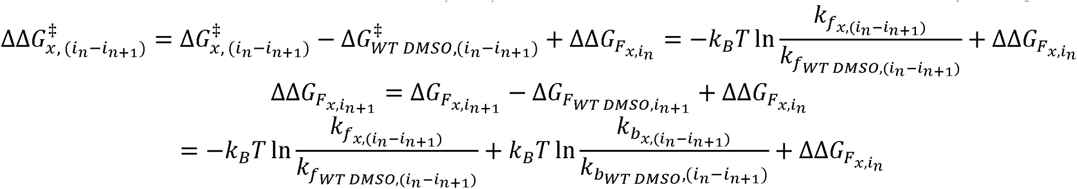

where *n* ranges from 1 to 6, with the corresponding intermediate states: *i*_1_ = 114 nm, *i*_2_ = 81 nm, *i*_3_ = 63 nm, *i*_4_ = 48 nm, *i*_5_ = 33 nm, and *i*_6_ = 21 nm. *x* represents the condition being examined. For *n* = 1, 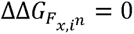, as the energy of the *U*_h_ state (114 nm) is equivalent across all the conditions, including WT and ΔF508 CFTR. *k*_f_ and *k*_b_ represent the forward and backward transition rates respectively. Both ΔΔ*G*^‡^ and ΔΔ*G*_F_ are expressed in units of *k*_B_*T*.

## Author Contributions

J.C. and T.-Y.Y. conceived of and supervised the project. S.A.K., J.L., J.C. and T.-Y.Y. designed the experiments. S.A.K. performed the single-molecule MT experiments. J.L. purified WT and mutant CFTR proteins. S.A.K. and T.-Y.Y. analyzed and visualized the MT assay data with inputs from other authors. S.A.K., J.L., J.C. and T.-Y.Y. wrote the manuscript.

## Acknowledgments

We thank Hak Chan Kim, Chanwoo Lee, and Hyun-Kyu Choi for their discussion in single-molecule MT data analysis, and Masa Predin for help with construct cloning. This work was supported by National Research Foundation of Korea (NRF) grant funded by the Korea government (MSIT) (RS-2021-NR059913 to T.-Y.Y.) and the Howard Hughes Medical Institute (to J.C.).

## Data availability

All data that support the findings of this study are available in the manuscript or and SI Appendix. Raw data have been deposited in Github (https://github.com/tyyoonlab-snu/Magnetic-Tweezers-2024/new/main).

## References

1. J. R. Riordan et al., Identification of the cystic fibrosis gene: cloning and characterization of complementary DNA. Science 245, 1066–1073 (1989).

2. C. E. Bear et al., Purification and functional reconstitution of the cystic fibrosis transmembrane conductance regulator (CFTR). Cell 68, 809–818 (1992).

3. M. P. Anderson et al., Nucleoside triphosphates are required to open the CFTR chloride channel. Cell 67, 775–784 (1991).

4. M. P. Anderson, D. P. Rich, R. J. Gregory, A. E. Smith, M. J. Welsh, Generation of cAMP-activated chloride currents by expression of CFTR. Science 251, 679–682 (1991).

5. M. L. Drumm et al., Correction of the cystic fibrosis defect in vitro by retrovirus-mediated gene transfer. Cell 62, 1227–1233 (1990).

6. Z. Zhang, J. Chen, Atomic structure of the cystic fibrosis transmembrane conductance regulator. Cell 167, 1586–1597. e1589 (2016).

7. F. Seibert et al., Influence of phosphorylation by protein kinase A on CFTR at the cell surface and endoplasmic reticulum. Biochimica et Biophysica Acta (BBA)-Biomembranes 1461, 275–283 (1999).

8. S. H. Cheng et al., Defective intracellular transport and processing of CFTR is the molecular basis of most cystic fibrosis. Cell 63, 827–834 (1990).

9. D. B. Sanders, A. Fink, Background and epidemiology. Pediatric Clinics of North America 63, 567 (2016).

10. D. C. Gadsby, P. Vergani, L. Csanády, The ABC protein turned chloride channel whose failure causes cystic fibrosis. Nature 440, 477–483 (2006).

11. J. Zielenski, L.-C. Tsui, Cystic fibrosis: genotypic and phenotypic variations. Annual review of genetics 29, 777–807 (1995).

12. S. M. Rowe, S. Miller, E. J. Sorscher, Mechanisms of disease: cystic fibrosis. New England Journal of Medicine 352, 1992–2001 (2005).

13. M. D. Amaral, C. M. Farinha, Rescuing mutant CFTR: a multi-task approach to a better outcome in treating cystic fibrosis. Current pharmaceutical design 19, 3497–3508 (2013).

14. J. R. Riordan, CFTR function and prospects for therapy. Annu Rev Biochem 77, 701–726 (2008).

15. R. R. Kopito, Biosynthesis and degradation of CFTR. Physiological reviews 79, S167–S173 (1999).

16. P. H. Thibodeau et al., The cystic fibrosis-causing mutation ΔF508 affects multiple steps in cystic fibrosis transmembrane conductance regulator biogenesis. Journal of Biological Chemistry 285, 35825–35835 (2010).

17. W. M. Rabeh et al., Correction of both NBD1 energetics and domain interface is required to restore ΔF508 CFTR folding and function. Cell 148, 150–163 (2012).

18. A. Zaher et al., A review of Trikafta: triple cystic fibrosis transmembrane conductance regulator (CFTR) modulator therapy. Cureus 13 (2021).

19. T. Okiyoneda et al., Mechanism-based corrector combination restores ΔF508-CFTR folding and function. Nature chemical biology 9, 444–454 (2013).

20. G. Veit, et al., Allosteric folding correction of F508del and rare CFTR mutants by elexacaftor-tezacaftor-ivacaftor (Trikafta) combination. JCI insight 5 (2020).

21. S. M. Rowe, A. S. Verkman, Cystic fibrosis transmembrane regulator correctors and potentiators. Cold Spring Harbor perspectives in medicine 3, a009761 (2013).

22. P. G. Middleton et al., Elexacaftor-tezacaftor-ivacaftor for cystic fibrosis with a single Phe508del allele. New England Journal of Medicine 381, 1809–1819 (2019).

23. J. L. Taylor-Cousar, P. D. Robinson, M. Shteinberg, D. G. Downey, CFTR modulator therapy: transforming the landscape of clinical care in cystic fibrosis. The Lancet 402, 1171–1184 (2023).

24. B. S. Quon, S. M. Rowe, New and emerging targeted therapies for cystic fibrosis. Bmj 352 (2016).

25. B. Kleizen, T. van Vlijmen, H. R. de Jonge, I. Braakman, Folding of CFTR is predominantly cotranslational. Molecular cell 20, 277–287 (2005).

26. B. Kleizen et al., Co-Translational Folding of the First Transmembrane Domain of ABC-Transporter CFTR is Supported by Assembly with the First Cytosolic Domain. J Mol Biol 433, 166955 (2021).

27. H. Lewis et al., Structure and dynamics of NBD1 from CFTR characterized using crystallography and hydrogen/deuterium exchange mass spectrometry. Journal of molecular biology 396, 406–430 (2010).

28. N. Soya, A. Roldan, G. L. Lukacs, Differential scanning fluorimetry and hydrogen deuterium exchange mass spectrometry to monitor the conformational dynamics of NBD1 in cystic fibrosis. Protein Misfolding Diseases: Methods and Protocols, 53–67 (2019).

29. C. L. Ward, R. R. Kopito, Intracellular turnover of cystic fibrosis transmembrane conductance regulator. Inefficient processing and rapid degradation of wild-type and mutant proteins. Journal of Biological Chemistry 269, 25710–25718 (1994).

30. N. Soya et al., Folding correctors can restore CFTR posttranslational folding landscape by allosteric domain-domain coupling. Nat Commun 14, 6868 (2023).

31. P. van der Sluijs, H. Hoelen, A. Schmidt, I. Braakman, The Folding Pathway of ABC Transporter CFTR: Effective and Robust. J Mol Biol 10.1016/j.jmb.2024.168591, 168591 (2024).

32. F. Liu, Z. Zhang, L. Csanady, D. C. Gadsby, J. Chen, Molecular Structure of the Human CFTR Ion Channel. Cell 169, 85–95 e88 (2017).

33. Z. Zhang, F. Liu, J. Chen, Molecular structure of the ATP-bound, phosphorylated human CFTR. Proc Natl Acad Sci U S A 115, 12757–12762 (2018).

34. Z. Zhang, F. Liu, J. Chen, Conformational Changes of CFTR upon Phosphorylation and ATP Binding. Cell 170, 483–491 e488 (2017).

35. J. Levring et al., CFTR function, pathology and pharmacology at single-molecule resolution. Nature 616, 606–614 (2023).

36. K. Fiedorczuk, J. Chen, Molecular structures reveal synergistic rescue of Delta508 CFTR by Trikafta modulators. Science 378, 284–290 (2022).

37. K. Fiedorczuk, J. Chen, Mechanism of CFTR correction by type I folding correctors. Cell 185, 158–168 e111 (2022).

38. F. Liu et al., Structural identification of a hotspot on CFTR for potentiation. Science 364, 1184–1188 (2019).

39. K. Varga et al., Efficient intracellular processing of the endogenous cystic fibrosis transmembrane conductance regulator in epithelial cell lines. Journal of Biological Chemistry 279, 22578–22584 (2004).

40. C. M. Farinha, S. Canato, From the endoplasmic reticulum to the plasma membrane: mechanisms of CFTR folding and trafficking. Cellular and molecular life sciences 74, 39–55 (2017).

41. H. K. Choi et al., Watching helical membrane proteins fold reveals a common N-to-C-terminal folding pathway. Science 366, 1150–1156 (2019).

42. H. K. Choi et al., Evolutionary balance between foldability and functionality of a glucose transporter. Nat Chem Biol 18, 713–723 (2022).

43. A. H. Keeble et al., Approaching infinite affinity through engineering of peptide–protein interaction. Proceedings of the National Academy of Sciences 116, 26523–26533 (2019).

44. D. Min, M. A. Arbing, R. E. Jefferson, J. U. Bowie, A simple DNA handle attachment method for single molecule mechanical manipulation experiments. Protein Sci 25, 1535–1544 (2016).

45. H. K. Choi, H. G. Kim, M. J. Shon, T. Y. Yoon, High-Resolution Single-Molecule Magnetic Tweezers. Annu Rev Biochem 91, 33–59 (2022).

46. L. Aleksandrov, A. A. Aleksandrov, X.-b. Chang, J. R. Riordan, The first nucleotide binding domain of cystic fibrosis transmembrane conductance regulator is a site of stable nucleotide interaction, whereas the second is a site of rapid turnover. Journal of Biological Chemistry 277, 15419–15425 (2002).

47. Y. Zhang, J. Jiao, A. A. Rebane, Hidden Markov Modeling with Detailed Balance and Its Application to Single Protein Folding. Biophys J 111, 2110–2124 (2016).

48. T.-H. Lee, Extracting kinetics information from single-molecule fluorescence resonance energy transfer data using hidden Markov models. The Journal of Physical Chemistry B 113, 11535–11542 (2009).

49. G. Lukacs et al., The delta F508 mutation decreases the stability of cystic fibrosis transmembrane conductance regulator in the plasma membrane. Determination of functional half-lives on transfected cells. Journal of Biological Chemistry 268, 21592–21598 (1993).

50. W. Dalemans et al., Altered chloride ion channel kinetics associated with the ΔF508 cystic fibrosis mutation. Nature 354, 526–528 (1991).

51. D. N. Sheppard et al., Mutations in CFTR associated with mild-disease-form CI-channels with altered pore properties. Nature 362, 160–164 (1993).

52. G. L. Lukacs et al., Conformational maturation of CFTR but not its mutant counterpart (delta F508) occurs in the endoplasmic reticulum and requires ATP. The EMBO journal 13, 6076–6086 (1994).

53. K. Du, M. Sharma, G. L. Lukacs, The ΔF508 cystic fibrosis mutation impairs domain-domain interactions and arrests post-translational folding of CFTR. Nature structural & molecular biology 12, 17–25 (2005).

54. H. A. Lewis et al., Impact of the ΔF508 mutation in first nucleotide-binding domain of human cystic fibrosis transmembrane conductance regulator on domain folding and structure. Journal of Biological Chemistry 280, 1346–1353 (2005).

55. P. G. Middleton et al., Elexacaftor–tezacaftor–ivacaftor for cystic fibrosis with a single Phe508del allele. New England Journal of Medicine 381, 1809–1819 (2019).

56. M. F. Rosser, D. E. Grove, L. Chen, D. M. Cyr, Assembly and misassembly of cystic fibrosis transmembrane conductance regulator: folding defects caused by deletion of F508 occur before and after the calnexin-dependent association of membrane spanning domain (MSD) 1 and MSD2. Molecular biology of the cell 19, 4570–4579 (2008).

57. W. E. Balch, D. M. Roth, D. M. Hutt, Emergent properties of proteostasis in managing cystic fibrosis. Cold Spring Harbor perspectives in biology 3, a004499 (2011).

58. J. Im et al., ABC-transporter CFTR folds with high fidelity through a modular, stepwise pathway. Cell Mol Life Sci 80, 33 (2023).

59. C. B. Anfinsen, Principles that govern the folding of protein chains. Science 181, 223–230 (1973).

60. S. P. Cole, Multidrug resistance protein 1 (MRP1, ABCC1), a “multitasking” ATP-binding cassette (ABC) transporter. Journal of Biological Chemistry 289, 30880–30888 (2014).

61. R. M. Vernon et al., Stabilization of a nucleotide-binding domain of the cystic fibrosis transmembrane conductance regulator yields insight into disease-causing mutations. Journal of Biological Chemistry 292, 14147–14164 (2017).

62. T. Hillenaar, J. Beekman, P. van der Sluijs, I. Braakman, Redefining Hypo- and Hyper-Responding Phenotypes of CFTR Mutants for Understanding and Therapy. Int J Mol Sci 23 (2022).

63. L. Cui et al., Domain interdependence in the biosynthetic assembly of CFTR. Journal of molecular biology 365, 981–994 (2007).

64. K. Liu, X. Chen, C. M. Kaiser, Energetic dependencies dictate folding mechanism in a complex protein. Proceedings of the National Academy of Sciences 116, 25641–25648 (2019).

65. A. Khushoo, Z. Yang, A. E. Johnson, W. R. Skach, Ligand-driven vectorial folding of ribosome-bound human CFTR NBD1. Mol Cell 41, 682–692 (2011).

66. A. Goehring et al., Screening and large-scale expression of membrane proteins in mammalian cells for structural studies. Nature protocols 9, 2574–2585 (2014).

