## Supplementary Figures for "Single-molecule dissection of CFTR folding defects and pharmacological rescue"

**This PDF file includes:**

Figures S1 to S12

SI References

**Figures and Figure legends**

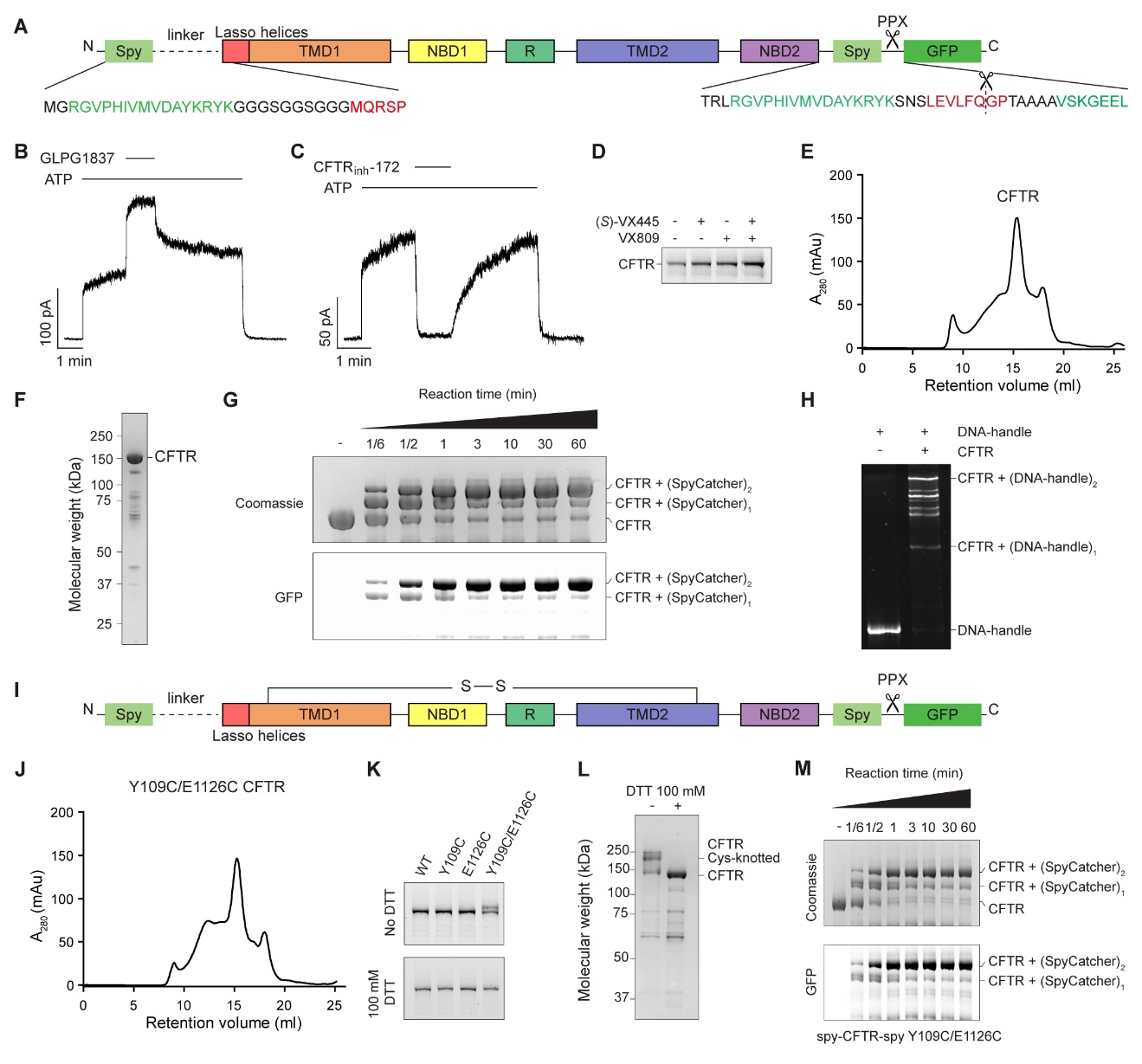

**Figure S1. Characterization and purification of the dual-SpyTagged CFTR variants.**

(A) Schematic representation of a dual-SpyTagged CFTR variant containing a C-terminal PPX-cleavable GFP tag for purification. SpyTag is linked to the protein at both termini, allowing conjugation with SpyCatcher-DNA handles. (B and C) Representative recordings of GLPG1837-stimulated (B), and CFTR_inh_-172-inhibited (C) currents from dual-SpyTagged CFTR in inside-out excised patches. 3 mM ATP, 10 µM GLPG1837, and 10 µM CFTR_inh_-172 were used. (D) Dual-SpyTagged CFTR was transiently expressed in HEK293S GnTI^-^ cells with or without 1 µM VX809 (lumacaftor) and/or 0.2 µM (*S*)-VX445 (elexacaftor). Clarified detergent extracts were separated by SDS-PAGE under reducing condition and the gel was imaged for GFP fluorescence. (E) Gel filtration profile of digitonin-solubilized and purified dual-SpyTagged CFTR. (F) SDS-PAGE analysis of purified dual-SpyTagged CFTR. (G) Time-dependent conjugation of GFP-fused SpyCatcher to dual-SpyTagged CFTR. A mixture of 2 µM CFTR and 20 µM SpyCatcher-GFP was incubated at room temperature. Formation of the conjugated products was visualized as gel-shifts in SDS-PAGE. The gel was imaged for GFP fluorescence and then Coomassie-stained. (H) SDS-PAGE analysis showing conjugation of SpyCatcher-DNA handles with dual-SpyTagged CFTR. 200 nM SpyCatcher-DNA handles and 2 µM CFTR in bicelle buffer were incubated at room temperature for 90 minutes before gel separation. The gel was SYBR safe-stained. (I) Schematic representation of a Cys-knotted CFTR variant restricting the unfolding to the lasso helices and NBD2. (J) Gel filtration profile of digitonin-solubilized and purified Cys-knotted CFTR. (K) Disulfide-dependent gel-shift. GFP-fused CFTR variants with one or two cysteines introduced by substitution were transiently expressed in HEK293S GnTI^-^ cells. Clarified detergent extracts were separated by SDS-PAGE under both reducing and non-reducing conditions and the gel was imaged for GFP fluorescence. A gel shift was observed under non-reducing conditions only for the Y109C/E1126C pair, indicating successful disulfide formation. (L) Gel-shift analysis with the purified dual-SpyTagged Cys-knotted CFTR. Disulfide formation was verified by a migration shift in SDS-PAGE under non-reducing conditions. The gel was Coomassie-stained. (M) Time-dependent conjugation of GFP-fused SpyCatcher to dual-SpyTagged Cys-knotted CFTR. A mixture of 2 µM CFTR and 20 µM SpyCatcher-GFP was incubated at room temperature. Formation of the conjugated products was visualized as gel-shifts in SDS-PAGE. The gel was imaged for GFP fluorescence and then Coomassie-stained.

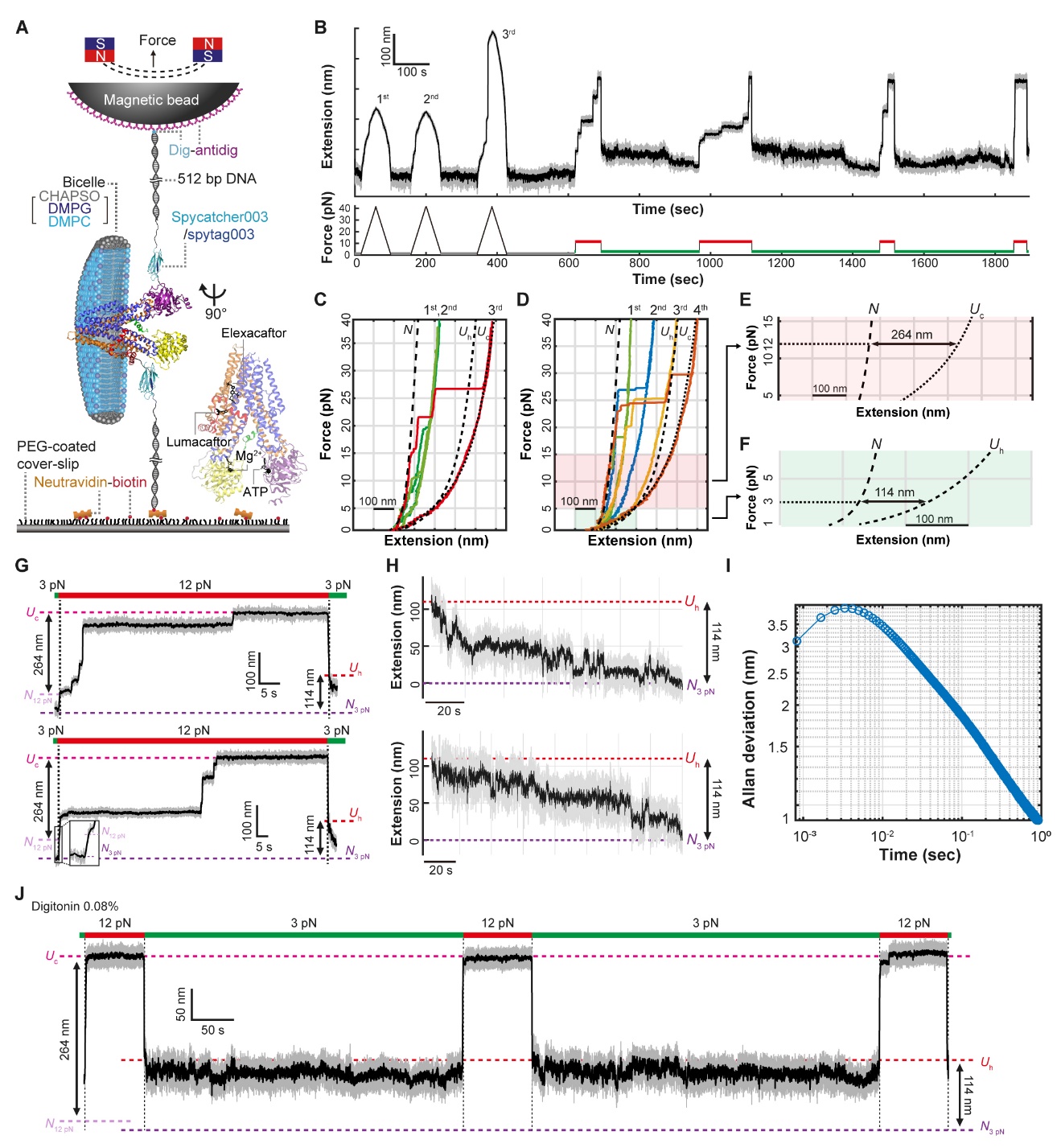

**Figure S2. Force cycles to observe the unfolding of a single CFTR protein.**

(A) Schematic representation of CFTR protein tethered to DNA handles in a single-molecule MT setup, allowing precise force application. The structure of CFTR (PDB: 5UAK) is shown, highlighting the positions where ligands bind (1-4). (B) Example of the force cycle used to monitor the unfolding and refolding of a single CFTR protein. After selecting a magnetic bead, several force-ramp cycles are conducted. After repeated force ramp cycles, the protein fully unfolds, with extension values aligning with theoretical predictions for CFTR's extension based on its structure. Following these force-ramp cycles, constant force cycles are employed to monitor unfolding and refolding events. (C) FECs from repeated force-ramp cycles in SI Appendix, Fig. S2B are shown in color, with theoretical FECs for the *N*, *U*_h_, and *U*_c_ depicted as black dotted lines for comparison. (D) Additional example of FECs showing the traversal of multiple unfolding intermediates, which are analyzed in detail in subsequent panels. (E) Enlarged view of the theoretical FECs for the *N* and *U*_c_ states, showing a theoretical extension of 264 nm at 12 pN. (F) Enlarged view of the theoretical FECs for the *N* and *U*_h_ states, showing a theoretical extension of 114 nm at 3 pN. (G) Representative unfolding traces obtained from constant force at 12 pN, illustrating individual CFTR proteins unfolded to the Uc state with a total unfolding extension of 264 nm. (H) Representative refolding traces obtained from constant force at 3 pN, illustrating individual CFTR proteins refolding from *U*_h_ to *N*, with a total extension of 114 nm. (I) Allan deviation of magnetic-bead fluctuation at 3 pN, measured using high-speed tracking at 1.2 kHz. When raw data is median filtered at 10 Hz (100 ms), the resolution is finer than 2 nm. (J) Representative trace showing CFTR behavior under detergent buffer.

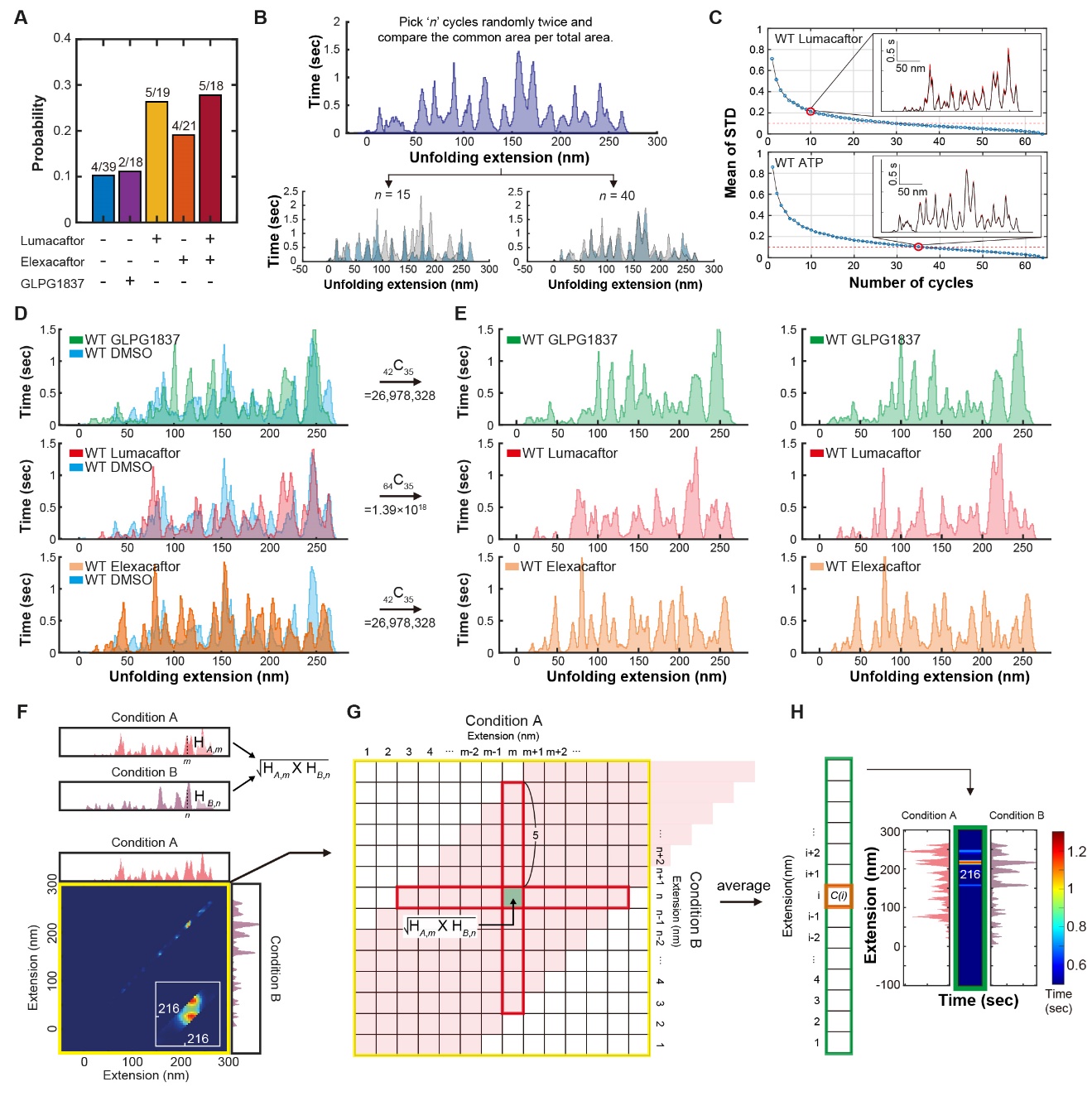

**Figure S3. Validation of single-molecule MT for probing CFTR unfolding and refolding.**

(A) Refolding probabilities for WT CFTR under the indicated conditions. The numbers above the bar graphs indicate the fraction of successfully refolded traces per total traces. 1 mM ATP, 10 µM GLPG1837, 1 µM lumacaftor and 1 µM elexacaftor was used. These concentrations were used for all subsequent MT experiments. (B and C) Convergence of unfolding histograms as the number of included cycles increases. When fewer than 30 cycles are used, histograms show notable variation (B, bottom left), but above 30 cycles, the histograms converge (B, bottom right). To quantify convergence, histograms were generated by randomly sampling increasing numbers of cycles (100 iterations per cycle number), and the mean standard deviation (STD) across all extension values was calculated (C). The inset shows that the STD is high when only 10 cycles are used, but markedly decreases with 35 cycles. (D) Unfolding histograms obtained under various ligand conditions in a blinded manner. (E) Two representative histograms generated by randomly selecting 35 cycles from the dataset in (D). The histograms on the right show these subsets, with the number below the arrows indicating the total number of possible 35-cycle combinations from the dataset. The similar outcomes of these histograms demonstrate the reproducibility of characteristic drug-specific unfolding patterns. (F–H) Cross-correlation analysis was performed to identify peaks that are prominent under conditions sharing a common ligand. Specifically, by multiplying the peak values from two histograms, shared features were emphasized, yielding high correlation values only when both conditions exhibited elevated peaks. For each pairwise comparison (e.g., Condition A vs. Condition B), a heatmap matrix was generated (F and G, yellow boxes), where each element represents the square root of the product of the corresponding histogram heights. To simplify the two-dimensional matrix, the data were averaged to generate a one-dimensional map (H, green box). In order to ensure accurate correlation calculation value at *i* (H, orange), we accounted for the widths of the peaks by including values within an 𝑚(𝑛)±5 nm range around each extension (0 < 𝑚(𝑛) < 264) (G, red). The detailed equation used to assess the correlation between two histograms is provided in the Methods.

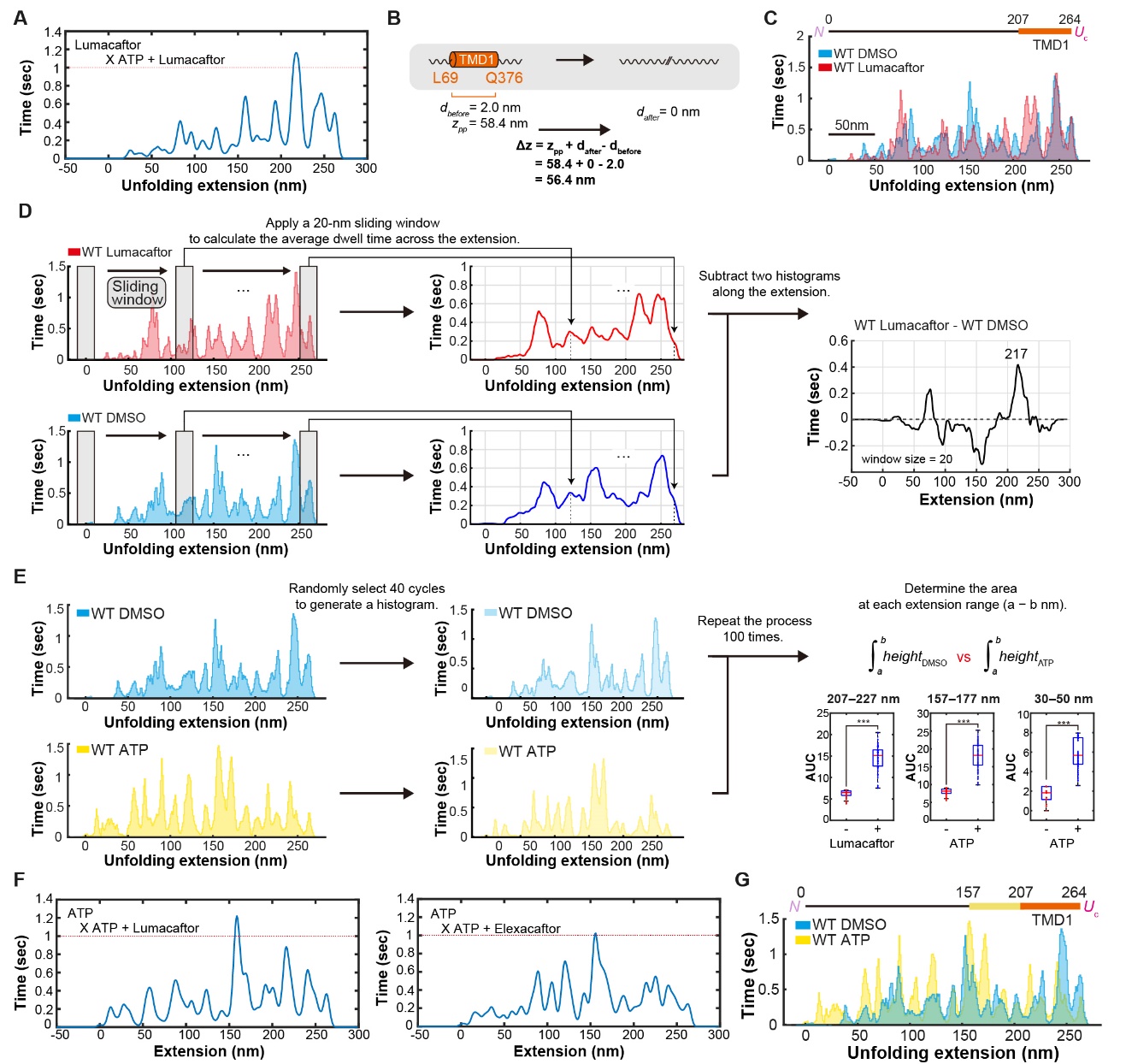

**Figure S4. Detailed unfolding analysis to understand the molecular identity from high-force unfolding.**

(A) The heatmap data from Fig. 1E is replotted as raw histogram data, illustrating cross-correlation results for WT Lumacaftor and WT ATP + lumacaftor. (B) Schematic representation of TMD1 unfolding and the theoretical calculation of extension change (Δ*z*) during the unfolding process. The contour length increase per unfolded domain (z_pp_) is estimated by multiplying the number of residues in the unfolded segment by the extension coefficient of a single amino acid under a 12 pN force with letters below marking the N- and C-terminal residues involved in the tertiary structure formation. Equation below the unidirectional arrow indicates an extension change associated with the domain's unfolding transition. The distance between the force application points 𝑑 was considered, including 𝑑*_before_* (pre-unfolding distance) and 𝑑*_after_* (post-unfolding distance). (C) Direct comparison of WT CFTR unfolding histograms with or without lumacaftor. *n* = 64 (WT Lumacaftor) and *n* = 45 (WT DMSO) force cycles were used to generate each histogram. (D) Schematic of sliding window difference analysis. A 20-nm sliding window was applied to dwell time histograms (left) to compute average dwell times along the unfolding extension (middle). The resulting histograms from two conditions were subtracted to identify regions with the greatest dwell time differences (right), highlighting structural segments most affected by experimental variation. (E) Methodology for AUC analysis to assess structural stabilization upon ligand binding. From the total set of unfolding cycles for each condition (left), 40 cycles, exceeding the minimum required for histogram convergence (see SI Appendix, Fig. S3C), were randomly selected to generate a histogram (middle). This process was repeated 100 times to calculate AUC values within defined extension ranges (denoted as 𝑎, 𝑏) (right). The boxplot elements represent the following: median (red vertical line); box limits (25^th^ and 75^th^ percentiles); whiskers (extending to 1.5×IQR). Statistical significance was tested using a two-sample t-test (***p ≤ 1 × 10^-3^). (F) The heatmap data from Fig. 1H is replotted as raw histogram data, illustrating cross-correlation results for WT ATP and WT ATP and a folding corrector. (G) Direct comparison of WT CFTR unfolding histograms with or without ATP. *n* = 64 (WT ATP) and *n* = 45 (WT DMSO) force cycles were used to generate each histogram.

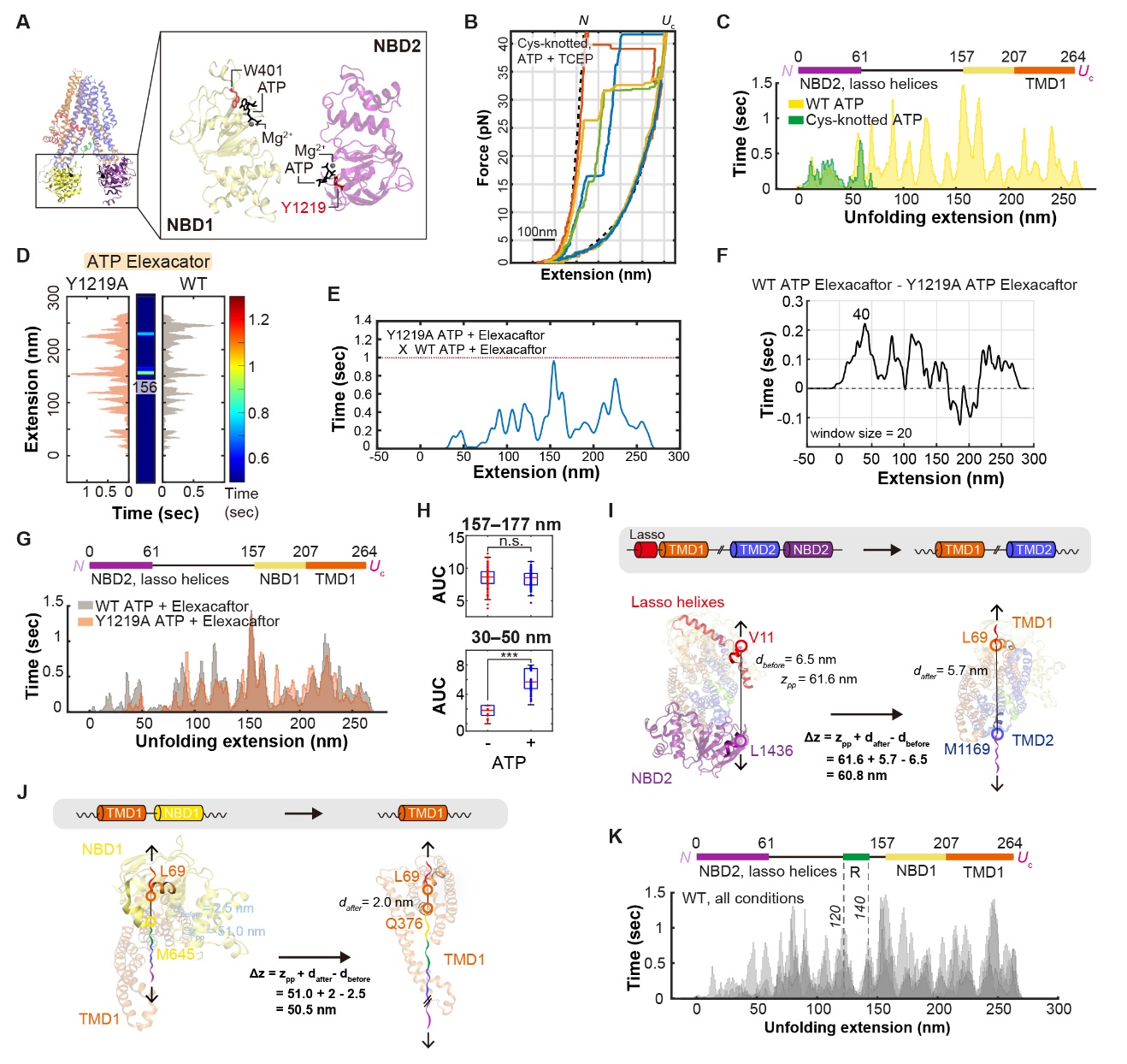

**Figure S5. High-force unfolding to determine molecular identities of extension values.**

(A) Structural representation of dephosphorylated CFTR with ATP and Mg²⁺ bound to both NBDs (PDB: 8FZQ) (left) (2). Enlarged view of ATP and Mg^2+^ bound to the NBDs (right). (B) Representative FECs of Cys-knotted CFTR in the presence of the reducing agent TCEP (colored traces). Upon reduction of the disulfide bond by TCEP, the fully unfolded Cys-knotted CFTR traces align with the *U*_c_ state FEC of WT CFTR, indicating full unfolding of all domains. (C) Direct comparison of WT CFTR unfolding histograms with or without lumacaftor. *n* = 45 (Cys-knotted ATP) and *n* = 64 (WT ATP) force cycles were used to generate each histogram. The WT ATP histogram is replotted from SI Appendix, Fig. S4G. (D) Cross-correlation analyses of Y1219A and WT CFTR unfolding histograms with ATP and Elexacaftor. The unfolding intermediate at 156 nm extension stabilized by ATP is annotated. *n* = 62 (Y1219A ATP + Elexacaftor) and *n* = 54 (WT ATP + Elexacaftor) force cycles were used to generate each histogram. The experiment was performed using 1 mM ATP and 1 µM elexacaftor. Both ATP and elexacaftor were included in order to solely isolate the effect of ATP binding to NBD2, independent of lasso helices folding, as both are expected to fold within a similar extension range. (E) The heatmap data from (D) is replotted as raw histogram data, illustrating cross-correlation results for Y1291A and WT with ATP and elexacaftor. (F) Sliding window difference analysis comparing unfolding histograms of Y1291A and WT with ATP and elexacaftor. A prominent increase in dwell time was observed between 30 and 50 nm, with the peak difference centered at 40 nm. *n* = 62 (Y1219A ATP and elexacaftor) and *n* = 54 (WT ATP and elexacaftor) force cycles were used to generate each histogram. (G) Direct comparison of Y1219A and WT CFTR unfolding histograms with ATP and elexacaftor. *n* = 62 (Y1219A ATP + Elexacaftor) and *n* = 54 (WT ATP + Elexacaftor) force cycles were used to generate each histogram. (H) Comparison of AUC in the 157–177 nm extension range (top) and 30–50 nm extension range (bottom). The boxplot elements represent the following: median (red vertical line); box limits (25^th^ and 75^th^ percentiles); whiskers (extending to 1.5×IQR). Statistical significance was tested using a two-sample t-test (***p ≤ 1 × 10^-3^). (I) Schematic illustrating the pulling geometry for NBD2 and lasso helices unfolding, resulting in 61 nm change in the extension. As the force is applied, the protein is assumed to undergo a rotational reconfiguration to accommodate unfolding. For clarity, the bicelle is omitted. (J) Schematic illustrating the pulling geometry for NBD1 unfolding, resulting in 51 nm change in the extension. (K) Overlaid unfolding histograms for WT CFTR treated with either DMSO, ATP, lumacaftor, or elexacaftor. A shared valley between 120 nm and 140 nm extension is highlighted.

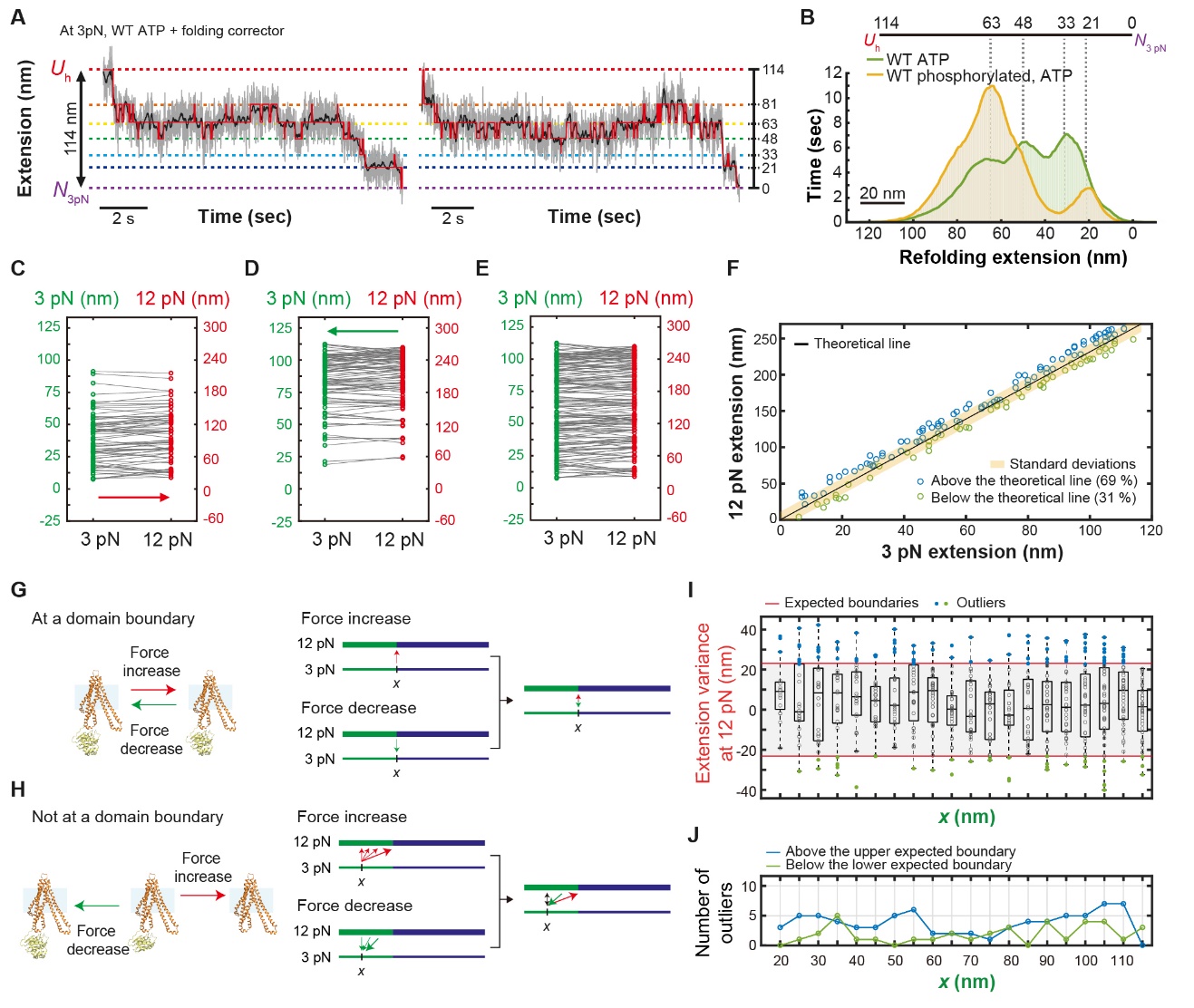

**Figure S6. Detailed analysis of CFTR folding dynamics.**

(A) Two representative folding traces in the presence of ATP and a folding corrector at 3 pN showing a total extension reduction of 114 nm. This is consistent with complete refolding from the *U*_h_ state to *N* state. 1.2 kHz raw data is gray, 10-Hz median filtered data is black, and the Viterbi path was overlaid on the data in red. Horizontal lines indicate extension values shown on the right. (B) Refolding histograms, generated with a 10-Hz median filter. The refolding histogram for WT ATP is replotted from Fig. 2. Peaks indicative of refolding intermediates are indicated with dashed lines. *n* = 34 (WT ATP) and *n* = 24 for (WT phosphorylated, ATP) refolding traces were used to generate each histogram. (C) Pairwise correlations of extension values measured at 3 pN and 12 pN immediately following a force jump from 3 pN to 12 pN (*n* = 72). (D) A similar analysis to (C), correlating extension values before and after decreasing the force from 12 pN to 3 pN (*n* = 92). (E) Aggregated correlations of extension values from all force-jump experiments, providing a comprehensive distribution of extension values across the full force range. (F) Theoretical line connecting (0,0) for the native state and (114, 264) for the fully unfolded state plotted with experimental pairwise extension values from force-jump experiments. Some extension values at 12 pN deviated from expectations, with more values falling above (blue circles) than below (green circles) the theoretical prediction. (G and H), Hypothetical model explaining variations in extension at 12 pN due to force-induced domain stability changes. At domain boundaries, minimal structural changes occur, resulting in a direct one-to-one correlation between extensions at 3 pN and 12 pN (G). Away from domain boundaries, additional unfolding upon force increase and partial refolding upon force decrease leads to higher-than-expected extension values at 12 pN (H). (I) Correlative mapping between 3-pN and 12-pN extension data (Fig. 2G) was analyzed by subtracting the median of the upper and lower expected domain boundaries. The top red line represents the expected boundary for *x*, while the bottom red line represents the expected boundary for *x*-20. Most data points fell within the domain boundaries (gray area), while outliers (blue and green dots) were observed, predominantly above these boundaries (blue dots). (J) Distribution of outliers relative to the expected boundaries shown in (I). Blue circles indicate outliers above the upper expected boundary (expected boundary for *x*), while green circles represent outliers below the lower expected boundary (expected boundary for *x*-20).

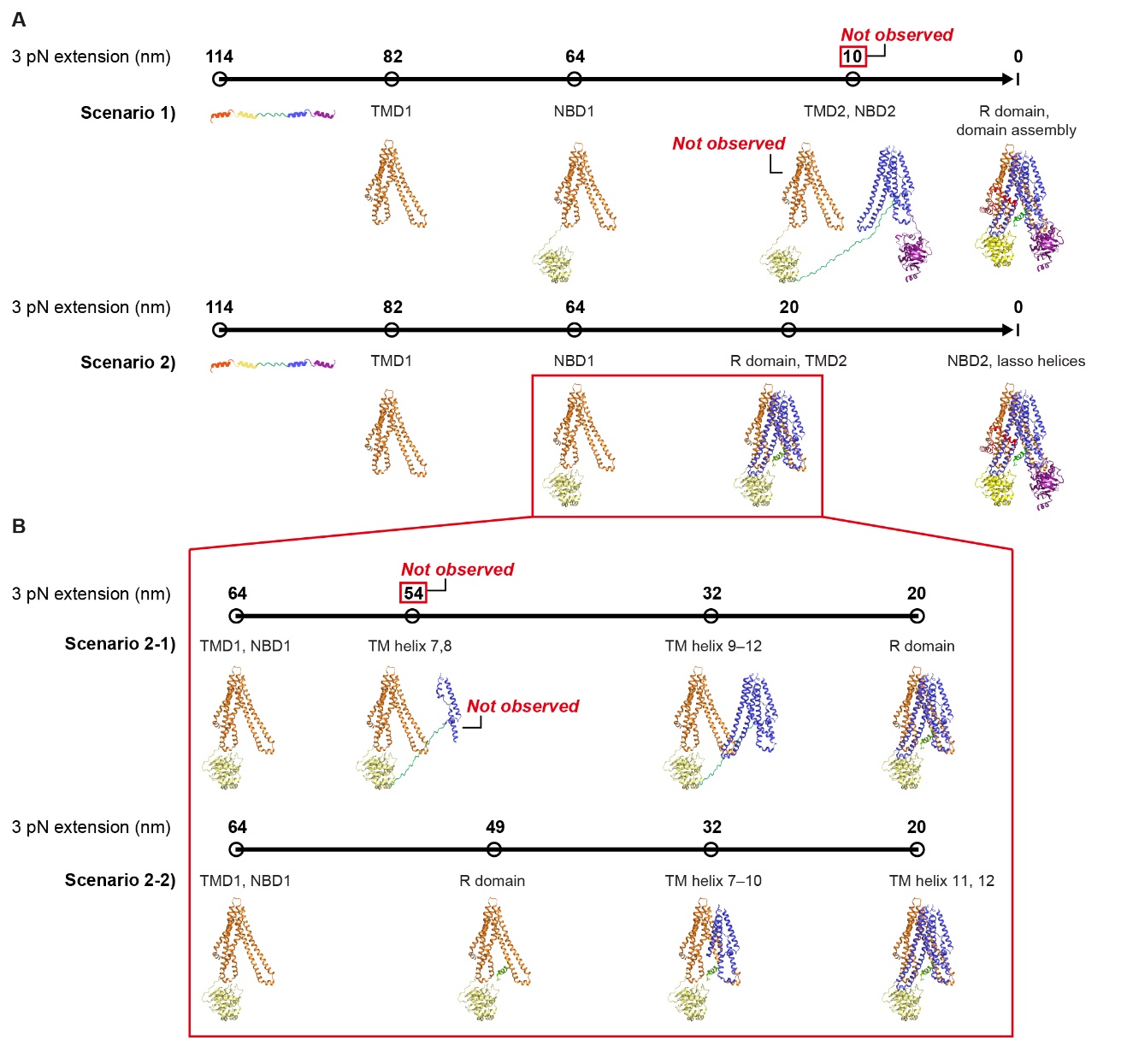

**Figure S7. Understanding the folding pathway of single CFTR.**

(A) Proposed CFTR folding models. Scenario 1: Independent domain folding. Each domain folds independently without requiring a pre-formed structure. However, this model is inconsistent with experimental data. While theoretical predictions suggest intermediates around 10 nm, the smallest experimentally observed intermediate during refolding is 21 nm, with no intermediates below 20 nm. Scenario 2: Sequential domain assembly. CFTR folds in a stepwise manner, where TMD2 assembles onto the pre-formed TMD1-NBD1 structure, followed by NBD2 folding. Numbers above the bold black lines indicate the theoretically expected extension values for each intermediate state while the domains labeled below correspond to the structural elements folded at each stage. (B) Submodels of Scenario 2. Scenario 2-1: Partial independent folding. TMD2 folds separately before assembling with the pre-formed TMD1-NBD1 structure, with R domain folding occurring during TMD2 assembly. Scenario 2-2: Templated folding. TMD2 folds directly onto the pre-assembled TMD1-NBD1 structure, with R domain folding preceding TMD2 assembly. Experimental data strongly support the templated folding model (Scenario 2-2). The consistent detection of a distinct intermediate at 48 nm across all conditions, prominently appearing in refolding histograms, suggests that this state corresponds to R domain folding. In contrast, the partial independent folding model predicts an alternative intermediate at 54 nm (corresponding to TM helices 7-8 folding), which was not observed experimentally. Numbers above the bold black lines indicate the theoretically expected extension values for each intermediate state while the domains labeled below correspond to the structural elements folded at each stage.

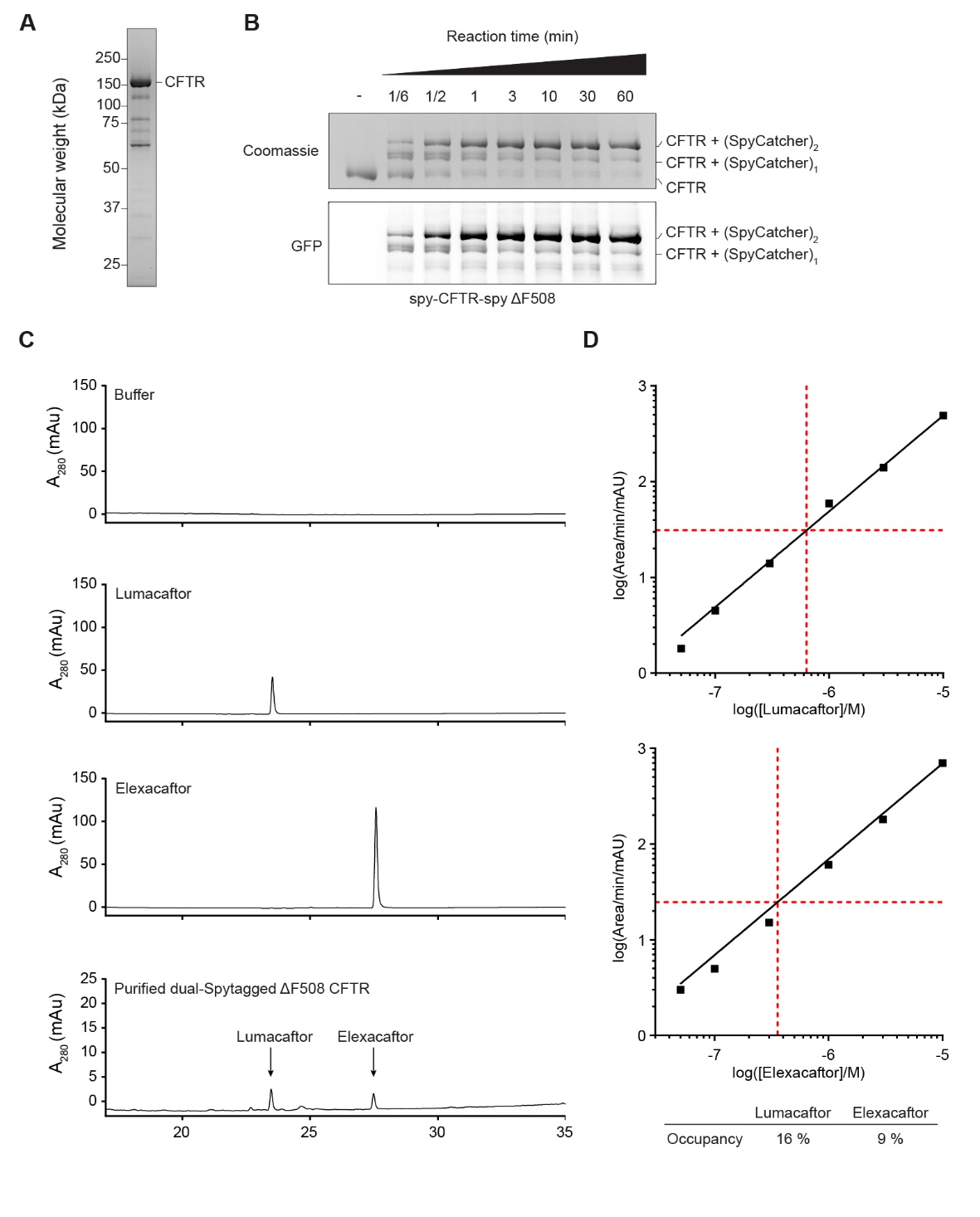

**Figure S8. Understanding the folding pathway of single CFTR.**

(A) SDS-PAGE analysis of purified ΔF508 dual-SpyTagged CFTR. (B) Time-dependent conjugation of GFP-fused SpyCatcher to dual-SpyTagged ΔF508 variant CFTR. A mixture of 2 µM CFTR and 20 µM SpyCatcher-GFP was incubated at room temperature. Formation of the conjugated products was visualized as gel-shifts in SDS-PAGE. The gel was imaged for GFP fluorescence and then Coomassie-stained. (C) HPLC profiles for buffer, pure lumacaftor, pure elexacaftor, and the purified and dialyzed ΔF508 CFTR sample. (D) Standards for the areas of the lumacaftor and elexacaftor HPLC peaks were generated by injection of samples of known concentration. Standard curves were generated by straight line fitting. The concentrations of lumacaftor and elexacaftor in the CFTR sample were estimated using these standards (indicated with red lines). The relative occupancies of lumacaftor and elexacaftor on ΔF508 CFTR were estimated to be 16 % and 9 %, respectively.

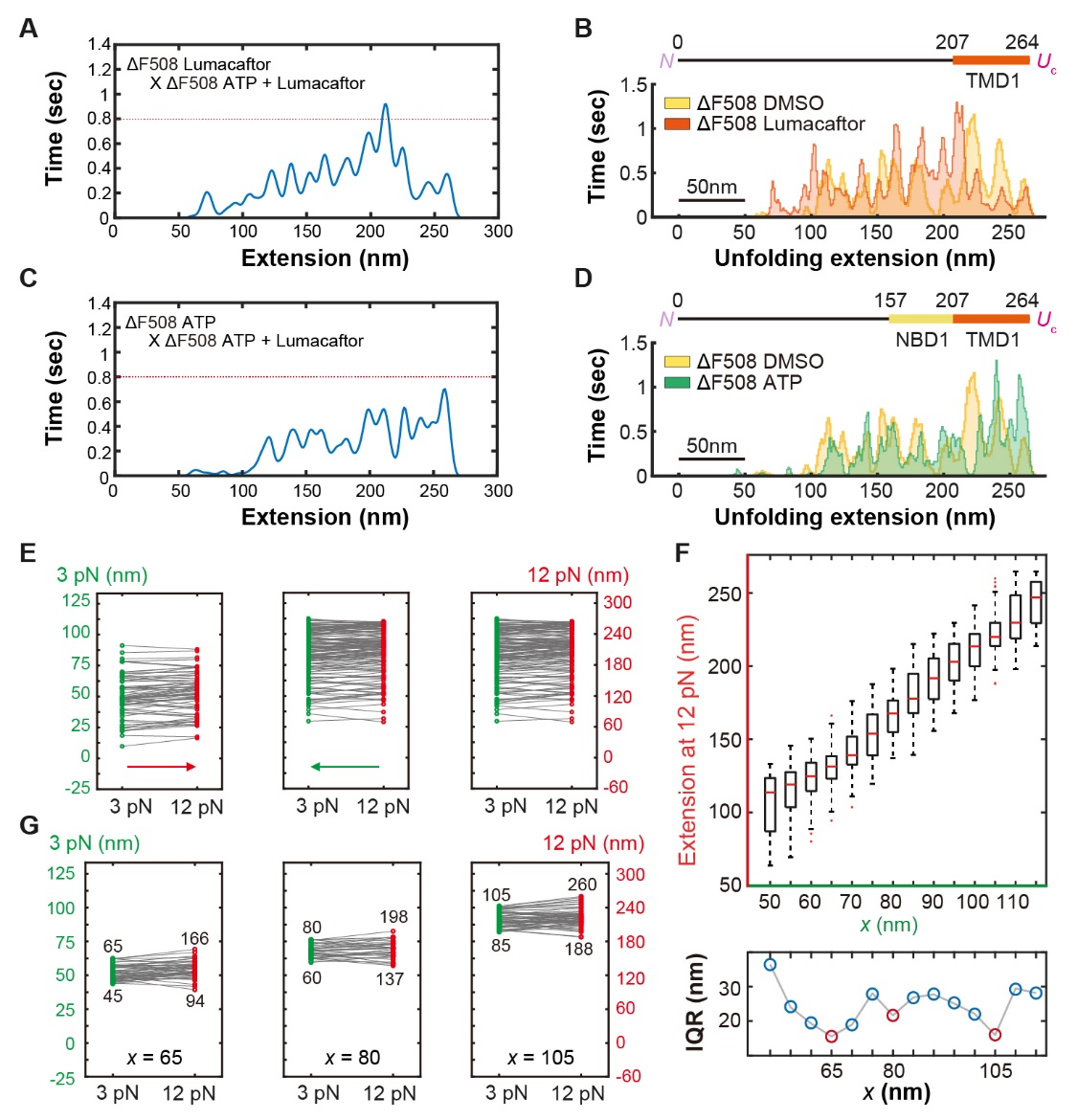

**Figure S9. Unfolding and refolding analyses to understand the folding pathway of ΔF508 CFTR.**

(A) The heatmap data from Fig. 3D is replotted as raw histogram data, illustrating cross-correlation results for ΔF508 Lumacaftor and ΔF508 ATP + Lumacaftor. (B) Direct comparison of ΔF508 CFTR unfolding histograms with or without lumacaftor. *n* = 70 (ΔF508 Lumacaftor) and *n* = 48 (ΔF508 DMSO) force cycles were used to generate each histogram. (C) The heatmap data from Fig. 3G is replotted as raw histogram data, illustrating cross-correlation results for ΔF508 ATP and ΔF508 ATP + Lumacaftor. (D) Direct comparison of ΔF508 CFTR unfolding histograms with or without ATP. *n* = 45 (ΔF508 ATP) and *n* = 48 (ΔF508 DMSO) force cycles were used to generate each histogram. (E) Pairwise correlations of extension values measured at 3 pN and 12 pN immediately following a force jump from 3 pN to 12 pN (*n* = 70) (left) and from 12 pN to 3 pN (*n* = 138) (middle). Aggregated correlations of extension values from all force-jump experiments, providing a comprehensive distribution of extension values across the full force range (right). (F) Correlative mapping between 3-pN and 12-pN extension spaces based on force-jump experiment data. Box plots depict the median (red line), IQR (box limits), and 1.5×IQR (whiskers). Data reflect *n* = 208 force-jump traces (top). IQR for different values of *x*. Local minima correspond to distinct structural regions at 65, 80 and 105 nm reflecting domain boundaries, indicating a structural transition point (bottom). (G) Mapping the exact extension values at 12 pN for the local minima identified in (F).

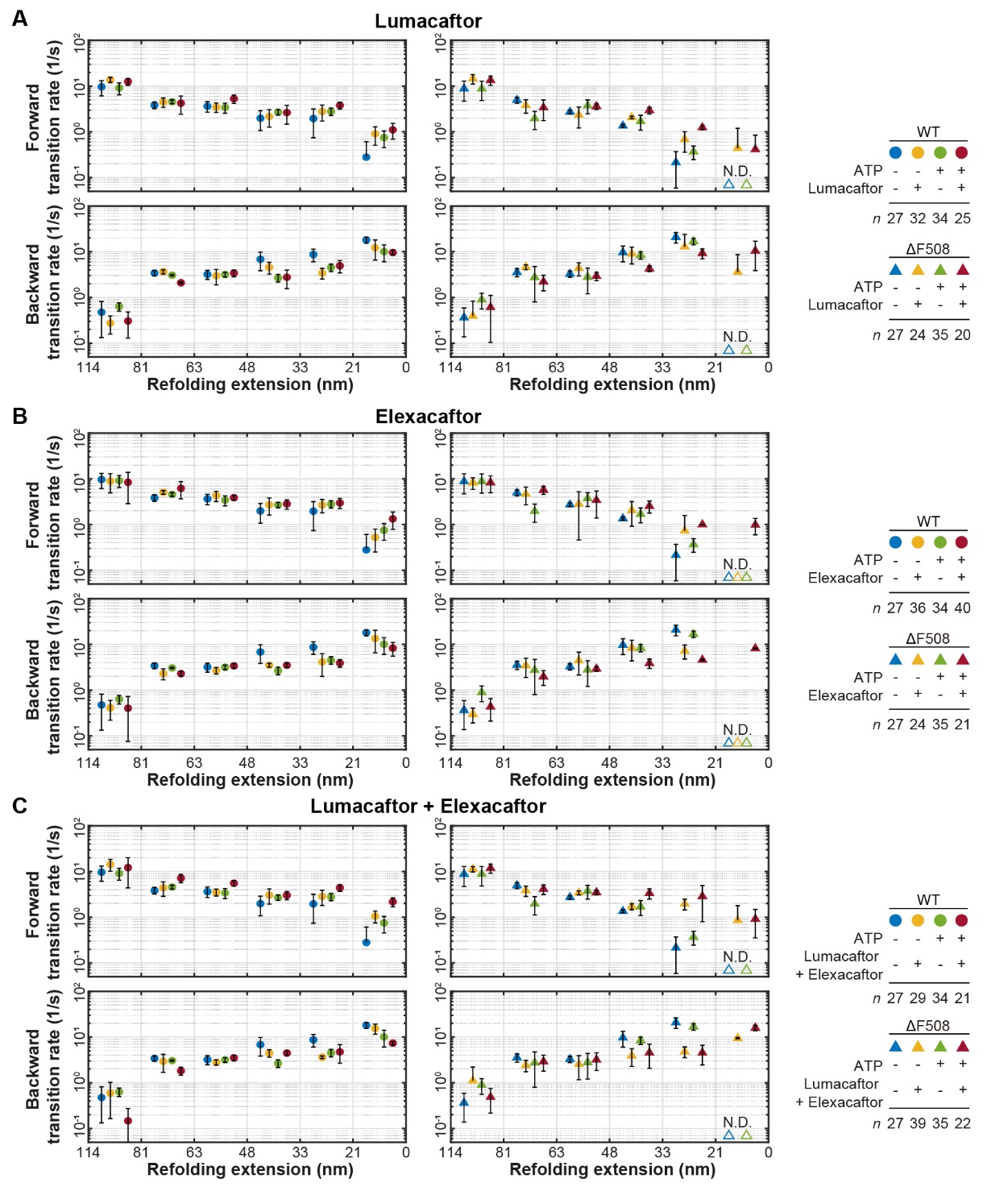

**Figure S10. Forward and backward rates for refolding transitions of WT and ΔF508 CFTR.**

(A–C), Forward and backward transition rates determined under the conditions indicated for WT and ΔF508 CFTR at 3 pN. The folding correctors used for individual experiments are lumacaftor (A), elexacaftor (B) and both lumacaftor and elexacaftor (C). Data represent means and standard deviations for *n* individual traces. In each of the plots, DMSO and ATP conditions are replotted for reference. N.D., not determined.

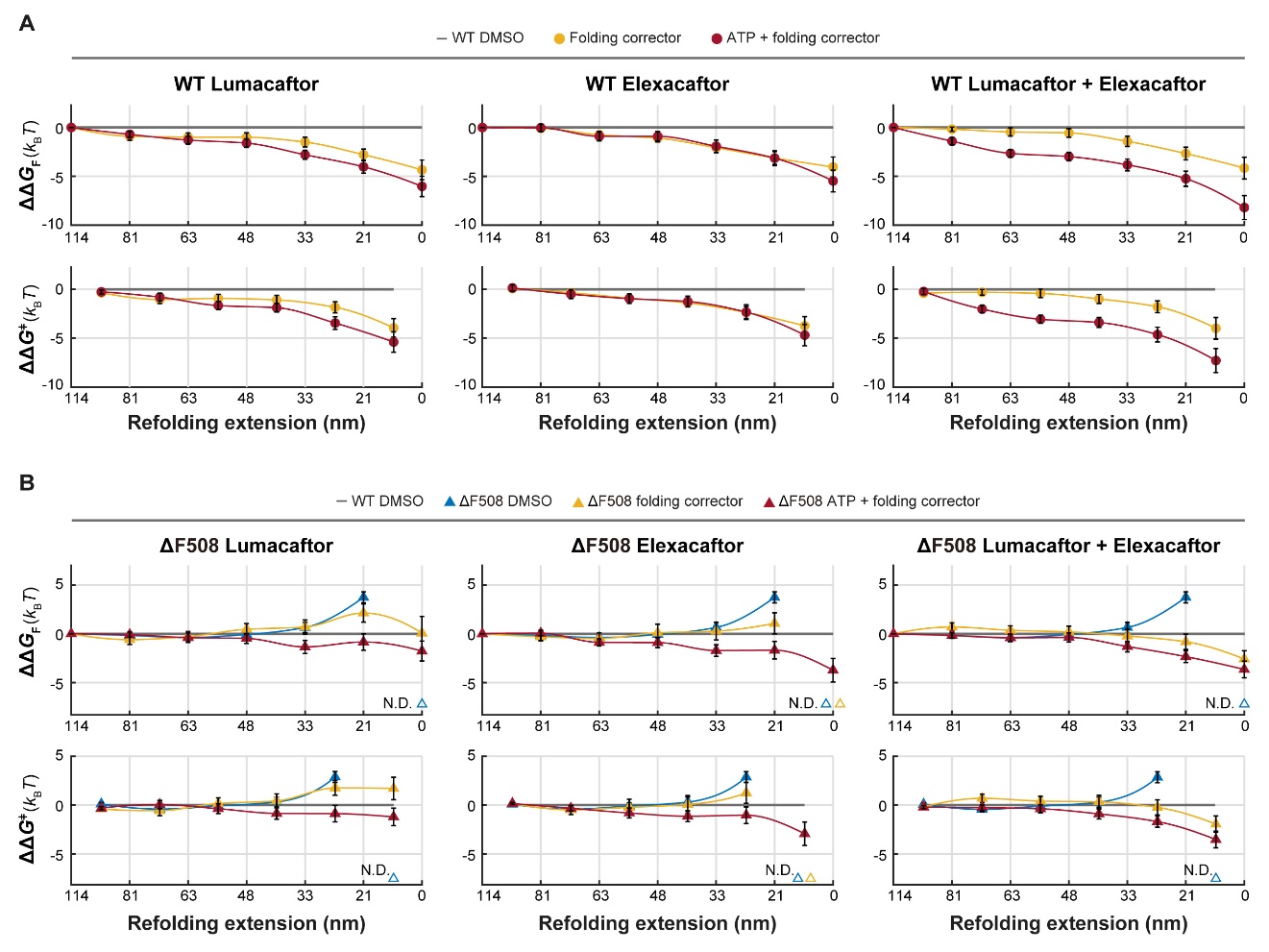

**Figure S11. ΔΔ*G*^‡^ and ΔΔ*G*_F_ analyses of WT and ΔF508 CFTR.**

(A and B) ΔΔ*G*_F_ and ΔΔ*G*^‡^ values for WT (A) or ΔF508 (B) CFTR for all transitions across various combinations of folding correctors and ATP. Values are calculated relative to WT DMSO (gray line at ΔΔ*G* = 0). Data represent means and propagated errors calculated from transition rate values obtained from *n* individual traces. ΔΔ*G* values are replotted from Fig. 5 B and C (ΔF508: DMSO; Lumacaftor + Elexacaftor) and Fig. 6D and E (WT: Lumacaftor + Elexacaftor; ATP + Lumacaftor + Elexacaftor, ΔF508: DMSO; Lumacaftor + Elexacaftor; ATP + Lumacaftor + Elexacaftor). N.D., not determined.

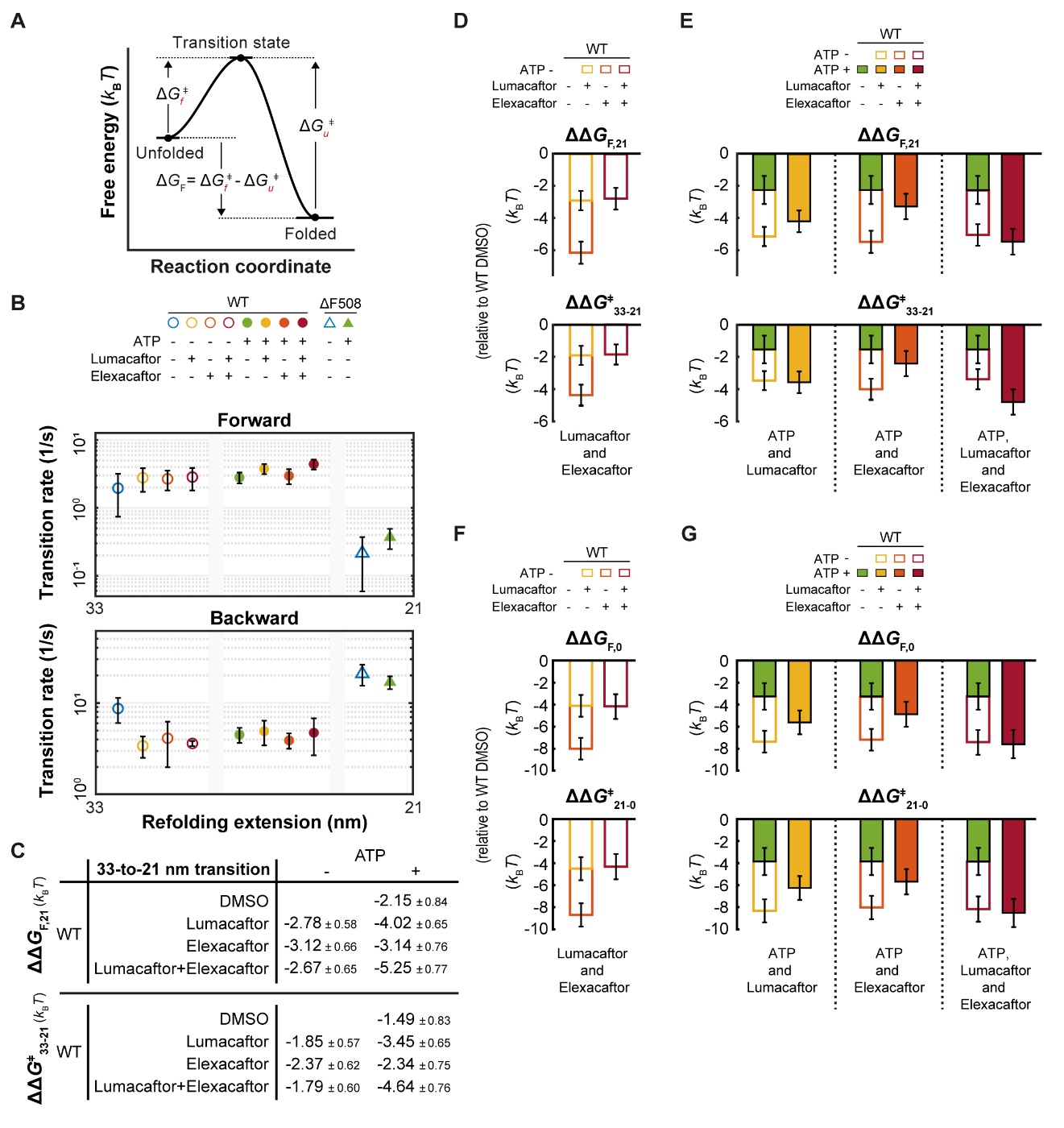

**Figure S12. Effects of folding correctors and ATP on WT and ΔF508 CFTR.**

(A) Schematic of energy barriers. Δ*G_f_*^‡^ and Δ*G_u_*^‡^ denote the free energy differences from the unfolded and folded states to the transition state, respectively. Δ*G_F_* is defined as Δ*G_f_*^‡^ - Δ*G_u_*^‡^. (B) Forward and backward transition rates at the 33-to-21 nm transition in WT and ΔF508 CFTR under various combinations of lumacaftor, elexacaftor, and ATP. Data represent means and standard deviations for *n* individual traces, with exact *n* values provided in SI Appendix, Fig. S10. (C) ΔΔ*G*_F,21_ and ΔΔ*G*^‡^_33-21_ values for all tested conditions in WT CFTR. Values for WT DMSO and WT ATP are replotted from Fig. 5E. (D) Comparative analysis of ΔΔ*G* values at 33-to-21 nm transition to evaluate the combinatorial effects of lumacaftor and elexacaftor in WT CFTR. ΔΔ*G* values are calculated relative to WT DMSO. Error bars represent propagated errors. (E) Comparative analyses of ΔΔ*G* values at 33-to-21 nm transition to evaluate the combinatorial effects of ATP and the folding correctors in WT CFTR. ΔΔ*G* values are calculated relative to WT DMSO. Error bars represent propagated errors. (F) Comparative analysis of ΔΔ*G* values at 21-to-0 nm transition to evaluate the combinatorial effects of lumacaftor and elexacaftor in WT CFTR. ΔΔ*G* values are calculated relative to WT DMSO. Error bars represent propagated errors. (G) Comparative analyses of ΔΔ*G* values at 21-to-0 nm transition to evaluate the combinatorial effects of ATP and the folding correctors in WT CFTR. ΔΔ*G* values are calculated relative to WT DMSO. Error bars represent propagated errors.

**SI References**

1.	F. Liu, Z. Zhang, L. Csanady, D. C. Gadsby, J. Chen, Molecular Structure of the Human CFTR Ion Channel. *Cell* **169**, 85-95 e88 (2017).

2.	J. Levring *et al.*, CFTR function, pathology and pharmacology at single-molecule resolution. *Nature* **616**, 606-614 (2023).

3.	K. Fiedorczuk, J. Chen, Molecular structures reveal synergistic rescue of Delta508 CFTR by Trikafta modulators. *Science* **378**, 284-290 (2022).

4.	K. Fiedorczuk, J. Chen, Mechanism of CFTR correction by type I folding correctors. *Cell* **185**, 158-168 e111 (2022).
